# Slow waves generation and propagation in a model of brain lesions

**DOI:** 10.1101/2025.11.12.688042

**Authors:** Gianluca Gaglioti, Leonardo Dalla Porta, Michele Angelo Colombo, Simone Russo, Thierry Nieus, Gustavo Deco, Maurizio Corbetta, Simone Sarasso, Maria V. Sanchez-Vives, Marcello Massimini

## Abstract

Slow waves (SWs), the hallmark of non-rapid eye movement (NREM) sleep, reflect the periodic occurrence of transient silent periods in cortical neurons (Down states). During NREM, SWs and Down states physiologically disrupt large-scale network interactions. Since early EEG studies, SWs have also been observed in awake patients after brain injury. Emerging evidence indicates that these intrusions of sleep-like activity interfere with ongoing network activity and contribute to motor and cognitive deficits; yet, the mechanisms governing the generation and spread of post-lesional SWs remain unclear. Here, we extend a neural mass model of EEG to capture transitions between wake-like and sleep-like dynamics and embed it in connectome-based networks with virtual lesions. This model supports that local disfacilitation, topology-dependent propagation, and synchrony-dependent amplification throughout the connectome are sufficient to produce post-lesional SWs. These mechanisms reproduce the spatial gradients of post-lesional SWs seen in patients and identify actionable targets for neuromodulation and rehabilitation.

## 1. Introduction

Brain lesions can affect remote areas beyond the site of structural injury^1–3^, disrupting network-level communication and impairing cognitive and behavioral function^4–7^, a phenomenon known as diaschisis^8^. Understanding the neuronal mechanisms underlying these network-wide effects remains a fundamental challenge in neurology, carrying important implications for rehabilitation and treatment^9^. In 1937, Grey Walter first described the appearance of slow waves (SWs) similar to those observed during NREM sleep in the brains of patients with brain lesions^10^. Recent evidence suggests that these SWs may play a crucial role in mediating the large-scale functional network alterations following focal brain injury^11^.

SWs emerge when a large neuronal population alternates synchronized periods of neuronal firing (Up states) and neuronal silence (Down states)^12–15^, a phenomenon also referred to as cortical bistability^16^. Mechanistically, Down states occur because of neuronal adaptation mediated by activity-dependent K^+^ currents^17^, as well as because of reductions in excitatory drive (disfacilitation)^18^ and/or inhibitory activity^19,20^. Although SWs are most prominent during NREM sleep, they can also occur locally during physiological wakefulness, especially after sleep deprivation^21–25^. SWs have also been observed in awake subjects following brain lesions, where they are most pronounced around the site of injury but can also extend to distant foci^10,26,27^.

The mechanisms of post lesional SWs are still elusive, and multiple factors may be at play^11^. Depending on lesion extent and location, disconnection may locally disrupt ascending activating inputs, enhancing activity-dependent adaptation mechanisms^28–30^, and/or disrupt lateral excitatory connections, causing disfacilitation and excitation/inhibition unbalance^19,31–34^. Once generated, SWs can then propagate across connected regions^30,35^, thus interfering with the functional dynamics of large-scale networks. SWs are known to disrupt network interactions and their intrusion in the brain of awake patients is clinically relevant^21,23,24^. Hence, understanding the mechanisms responsible for the generation and propagation of post-lesional SWs during wakefulness is essential to explain—and ultimately re-normalize—the functional consequences of brain injury.

In the present study, we tackle this question by adopting a multiscale computational approach. Large-scale computational models combine biologically informed network architecture from neuroimaging with node-level neural dynamics^36,37^, allowing us to examine how local interactions scale up to complex systems behavior in health and disease. This framework has already proven useful in epilepsy research^38^, where it is extensively used to study seizure initiation and propagation^39–41^.

Here, we develop a novel computational approach to study generation and propagation of SWs in cortical networks following brain injury. We adopt a Jansen-Rit (JR) model^42^, which has been extensively used to simulate spontaneous and evoked electroencephalography (EEG) and local field potential (LFP) oscillatory dynamics in awake-like conditions^43–48^. Since the original formulation lacks the adaptation dynamics required to reproduce SWs^17,49^, we extended the JR model by introducing an activity-dependent adaptation mechanism (Fig. 1A–C)^50^, enabling the model to capture the transition between wake-like and sleep-like states. Finally, by embedding this extended model into simplified motifs, topologically organized networks, and whole-brain connectomes, we systematically examined whether and how virtual lesions—mimicking either disconnection from ascending inputs or disruption of lateral cortico-cortical connectivity—give rise to perilesional SW generation and large-scale propagation (Fig. 1D).

**Figure 1.**
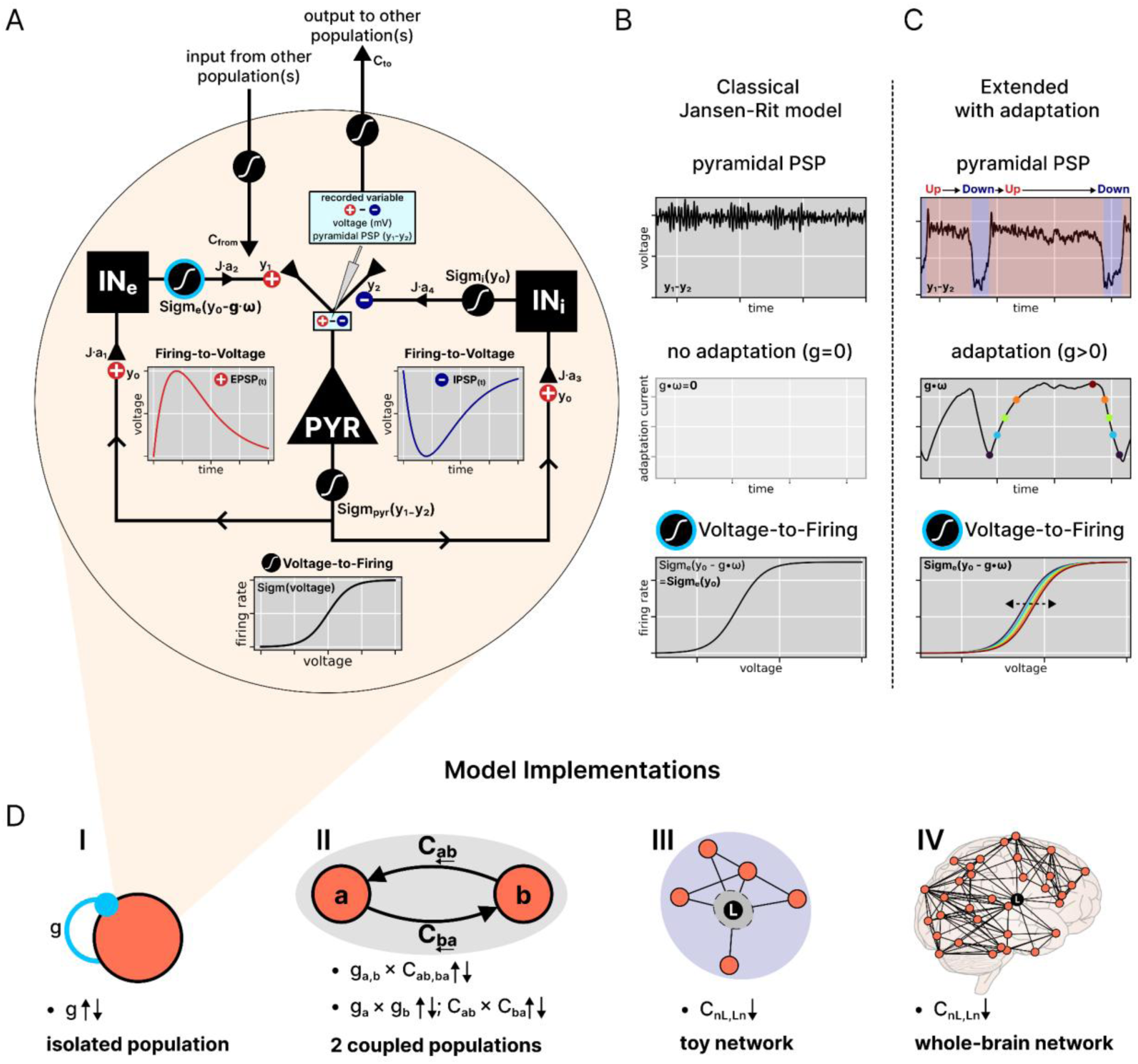
Extension of the neural mass model with spike-frequency adaptation mechanism across different scales: from a single-population to whole-brain network. (A) Representation of the Jansen-Rit (JR) model loop^62^, consisting of interconnected neuronal pools: pyramidal neurons (PYR), excitatory interneurons (IN_e_), and inhibitory interneurons (IN_i_). The model converts postsynaptic potentials back to firing rates via Voltage-to-Firing sigmoidal functions (Sigm) and transforms firing rates into postsynaptic potentials through Firing-to-Voltage operators. These Firing-to-Voltage operators differed for excitatory and inhibitory synapses (⊕ and ⊖, respectively). Excitatory postsynaptic potential (EPSP) generating functions are shown in red, while inhibitory postsynaptic potential (IPSP) generating functions are shown in blue. The IPSP is represented as negative for illustrative purposes to highlight its inhibitory effect on the pyramidal population. The recorded variable represents the pyramidal postsynaptic potential (PSP = y_1_ - y_2_), reflecting the net average postsynaptic potential from both excitatory and inhibitory inputs impinging on pyramidal neurons. This PSP also constitutes the output from the neural mass population to other populations, and is conventionally used as a proxy for EEG or LFP activity. Parameter definitions: J represents the average number of synapses between neuronal pools; a_1_, a_2_, a_3_, a_4_ are weight coefficients determining synaptic connection strength; y_0_, y_1_, y_2_ denote average postsynaptic membrane potentials of pyramidal, excitatory interneuron, and inhibitory interneuron populations, respectively. Notably, two parameters define the spike frequency adaptation, thereby extending the classical Rit-Jensen model: g is the constant that controls adaptation strength; ω represents the adaptation state-variable over time. Comparison between classical Jansen-Rit model (B) and extended version with adaptation (C). (B) The classical model generates standard pyramidal PSP oscillations without adaptation (coinciding with g=0). (C) Extended model with adaptation (g>0) exhibits Up and Down dynamics. The adaptation mechanism shifts the sigmoidal Voltage-to-Firing curve rightward during periods of high activity, reducing neuronal responsiveness and creating characteristic Up-to-Down state transitions observed in neural recordings; vice versa a leftward shift of the sigmoid increase sensitivity to input voltage during quiet periods, leading to Down-to-Up state transitions. (D) Schematic representations of the network scales explored in the study. I. *Isolated population*: single JR population model with variations in synaptic adaptation strength (g). II. *Two-coupled populations*: two coupled JR populations with symmetric and asymmetric variations in adaptation g and excitatory coupling C. In the symmetric case, the parameters were varied equally in a and b (i.e., g_a,b_ × C_ab,ba_). In the asymmetric case, the parameters were varied independently in the two populations (i.e., g_a_ × g_b_, and C*_ab_* × C*_ba_*). III. *Toy network*: simplified network exploring lesion effects. The lesion was implemented by disconnecting the lesion node (L) (i.e., setting to 0 all the incoming connections *C_Ln_,* and outgoing connections *C_nL_*). IV. *Whole-brain network*: large-scale model based on human structural connectivity data, investigating the effects of lesions on brain-wide dynamics. Lesions were implemented as in the toy network. Adaptation is implemented at all scales (I–IV). It is depicted as a self-loop (light blue circle) only in the isolated population panel for clarity, to highlight its activity-dependent nature.

## 2. Methods

### 2.1 Jansen-Rit model with spike-frequency adaptation

The Jansen-Rit (JR) model^42^ describes the temporal evolution of postsynaptic membrane potentials and firing rates of interconnected neuronal pools, representing pyramidal neurons (pyr), excitatory interneurons (IN_e_), and inhibitory interneurons (IN_i_) (Fig. 1A). In this study, we used the implementation of the JR model available in The Virtual Brain (TVB) simulator and extended it by introducing a spike-frequency adaptation mechanism^50^. The dynamics of the neuronal pools evolve according to the following set of differential equations:

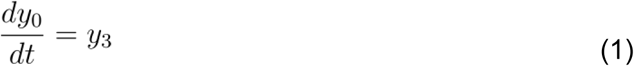

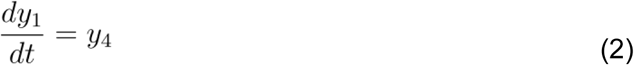

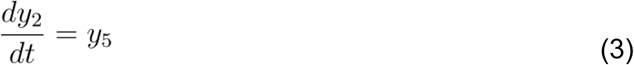

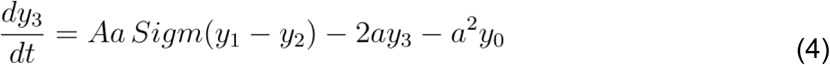

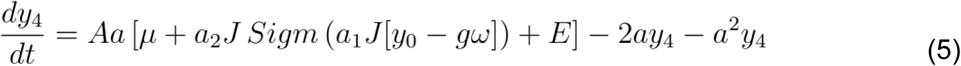

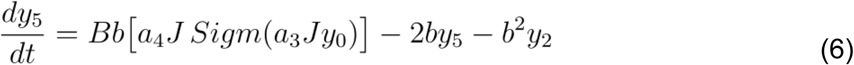

where y_0_, y_1_, and y_2_ represent the average postsynaptic membrane potentials (PSP) of pyr, IN_e_, and IN_i_, respectively. The corresponding firing rates are denoted as y_3_, y_4_, and y_5_. The parameters A and B represent the maximum amplitudes of excitatory and inhibitory postsynaptic potentials (EPSP and IPSP), while a and b are the inverse time constants governing synaptic dynamics. μ is the mean firing rate input to IN_e_ pool, J represents the average number of synapses between neuronal pools, and a_1_, a_2_, a_3_, a_4_ are weight coefficients determining the strength of synaptic connections.

Neuronal pools convert input membrane potentials into output firing rates through a sigmoidal activation function (Voltage-to-Firing, Fig. 1A):

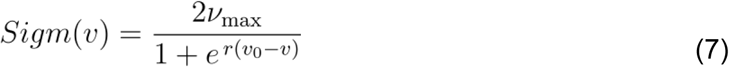

where ν_max_ is the maximum firing rate, v_0_ is the PSP threshold for which 50% firing rate is achieved and r determines the steepness of the sigmoidal function.

In Eq. 5, E represents the excitatory network input. The input to a given node *i* at time *t* is defined as the sum of the contributions from all nodes projecting to node *i*, expressed as:

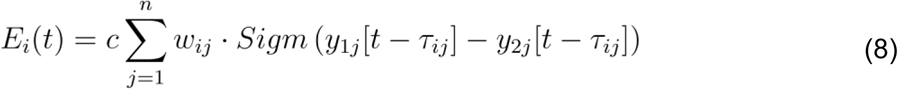

where w_ij_ is the connection weight from presynaptic node *j* to postsynaptic node *i*, and 𝜏_ij_ represents the time delay between *i* and *j*. The term c is the global coupling factor that scales all the connection weights. Thus, the effective excitatory coupling strength between *j* and *i* is given by the product of w_ij_ and c, and we denote it by C_ij_.

Spike-frequency adaptation (ω) is included as a slow dynamic process affecting the excitatory feedback loop:

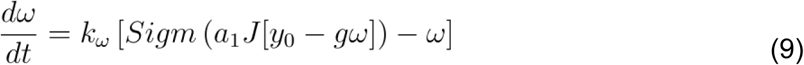

where *k_ω_* is the adaptation rate constant and *Sigm* is the sigmoidal input-output function of IN_e_ (Eqs. 5, 7; Fig. 1A-C). This formulation, proposed by Moran and collaborators, extends the classical Jansen-Rit model by incorporating an adaptation mechanism that reflects the activity-dependent reduction in neuronal excitability observed experimentally ^50^. This equation conforms to a universal phenomenological model^51^, which captures the essential dynamics of activity-dependent adaptation across a wide range of mechanisms and cell type: higher activity leads to a subsequent, relatively slow increase of the adaptation current. Increased adaptation shifts the system’s input-output (Voltage-to-Firing) curve to the right (Fig. 1C), requiring stronger depolarizing input to achieve the same firing rate in output. Due to the nonlinear sigmoidal shape, adaptation reduces the system’s responsiveness by decreasing the local gain (slope). Compared to the original formulation^50^ using only the adaptation variable *ω*, we introduced a scaling factor *g* that modulates its influence on the input–output function, thereby explicitly controlling the adaptation strength. When *g*=0, there is no adaptation, and the system reduces to the classical JR formulation (Fig. 1B).

Following the TVB convention, external noise is defined within the stochastic integration scheme and, in our case, it enters as an additive term in Eq. 5, simulating the spontaneous background activity at each node^42^. Noise was modelled as a white noise process with a standard deviation sd_noise_=10^-4^. The system was solved using the Heun stochastic method with a time step of 1 ms^42^, starting from random initial conditions and discarding the first 5 seconds of transient activity. The model parameters are listed in Supplementary Table 1.

The recorded variable in this study was the pyramidal postsynaptic potential (Fig. 1A), representing the net average postsynaptic potential, considering both excitatory and inhibitory input voltages impinging on pyramidal neurons (PSP = y_1_ - y_2_), which is conventionally used as a proxy for EEG or LFP activity in the source space^43,46,47^.

### 2.2 Isolated population model

A single JR population model (Fig. 1D-I) was implemented as a system of seven first-order ordinary differential equations (ODEs) as described by Eqs. 1-9 (excluding the network input). Bifurcation and fixed-point analyses of the deterministic JR model were performed using XPPAUT^52^. The bifurcation parameters were the average number of synapses between neuronal pools (J) and the adaptation strength (g), which were varied independently to assess changes in the qualitative behavior of the system. Local stability of fixed points was determined by XPPAUT via linearization around equilibria and computation of the eigenvalues of the Jacobian matrix. The stochastic version of the model was implemented in TVB and was used for all subsequent simulations of the study. The bifurcation diagram was constructed by collecting the 1st and 99th percentiles of PSP activity as a function of g. For each value of g, an independent simulation of 30s was performed, with g varied in steps of 1. The code used to run these analyses (in XPPAUT and TVB) is available on GitHub at https://github.com/gianlucagag/SWs-lesion_model

### 2.3 Two-population model

To assess the effects of cortical disconnection on the emergence of SWs, a two-population model was implemented (Fig. 1D-II). The model consisted of two JR populations, labeled JR*_a_* and JR*_b_*, which were coupled by bidirectional excitatory connections with no time delays. The system behavior was explored parametrically as a function of both symmetric and asymmetric variations of the adaptation strength (g) and the excitatory coupling strength (C) in the two populations, constructing 2D bifurcation diagrams in the parameter space. In the symmetric case, the parameters were varied equally in JR_a_ and JR_b_ (i.e., g vs C, where g_a_=g_b_, and C_ab_=C_ba_) (Fig 3 A). In the asymmetric case, the parameters were varied independently in the two populations (i.e., g_a_ vs g_b_, in Fig 3B and C_ab_ vs C_ba_ , where g_a_≠g_b_ and C_ab_≠C_ba_ in Fig 3D). Each simulation corresponding to a specific parameter configuration was run independently and lasted 30s. In these analyses, g was varied from 0 to 60 in steps of 0.5, while C was varied from 0 to 60 in steps of 1.

### 2.4 Toy network model

To relate the generation and propagation of SWs to simplified network motifs, a toy network model was implemented (Fig. 1D-III), incorporating delays, with physical distances and signal propagation velocity (v = 4 m/s) consistent with cortico-cortical connections ^53^. The network consisted of a perilesional subnetwork of five nodes (denoted Lₖ nodes, k=1..5) and a distant chain of three nodes (denoted Dₖ nodes, k=1..3). The connection weights w_i𝑗_ were normalized such that the sum of the in-weights of each node was equal to 1. With this normalization, the total excitatory input coupling to each node is determined by the global coupling factor c. To maintain the same level of excitability at each node, c was set to 30, as explored in the two-population model. The effective excitatory coupling between a presynaptic node *j* and a postsynaptic node *i* is therefore C_ij_ = cw_i𝑗_ (Eq. 8). In the perilesional network, each node had two afferent nodes, such that lesioning (disconnecting) any node resulted in a 50% reduction in the excitatory input to the connected Lₖ nodes. Lesions were simulated by setting all in- and out-weights of the lesioned node to zero. Twenty simulations were performed for each lesion condition and compared to a control condition (no lesion). Each simulation lasted 100s.

### 2.5 Human whole-brain network model

For the human brain network simulations, the model was implemented using structural brain connectivity data (Fig. 1D-IV). An open dataset was utilized, which provides brain structural connectivity matrices ready for modeling^54^. The dataset contains structural connectivity matrices from 88 healthy subjects. The matrices represent the connectivity among 90 cortical regions of interest (ROIs) as defined by Automatic Anatomical Labeling (AAL) ^55^. The 10 non-cortical regions were excluded since the JR model is designed for cortical networks only. Connectivity matrices were averaged across subjects, and only connections present in more than 50% of the subjects were retained, to obtain a sparse matrix retaining the typical within-subject density^56^. Subsequently, the weighted in-degree normalization was applied, and c=30 was set, as performed in the toy network. This normalization serves as a form of homeostatic regulation, equalizing the excitatory inputs received by the nodes while preserving the structural topology^44^. A control simulation was first run, followed by simulations in which each of the 80 nodes was lesioned (as in the toy network), and the effects on the remaining 79 nodes were analyzed. Each simulation lasted 100s.

### 2.6 Down state statistics

To detect *Down states* in the PSP, a fixed threshold (*th*) was used based on a model parameter representing the firing threshold (*v_0_*). *v_0_* (=5.52 mV) corresponds to the PSP value for which a 50% firing rate is achieved^42^. In other words, it is the input value in mV at the inflection point of the sigmoidal input-output function. The system was considered to be in a *Down state* whenever the PSP fell below *v_0_* (i.e. PSP<*th*), and in an *Up state* when the PSP exceeded it (i.e. PSP≥*th*). This simple threshold was chosen to effectively distinguish between low and high activity states in the simulation.

The %Down variable quantifies the percentage of time a node spends in the Down state. For a given node *i*, it was computed as the ratio of time spent in the *Down state* to the total simulation time, expressed as a percentage:

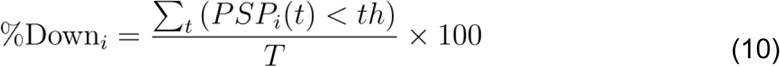

where PSP*_i_*(t) is the PSP of node *i* at time t, and T is the total simulation time.

The DownOverlap was computed as the percentage of time during which the PSP signals of two nodes simultaneously fell below the Down state threshold (*th*, Fig. 5C). For example, given nodes *i* and *j*, with corresponding signals *PSP_i_* and *PSP_j_*, the DownOverlap for node *i* relative to node *j* is defined as

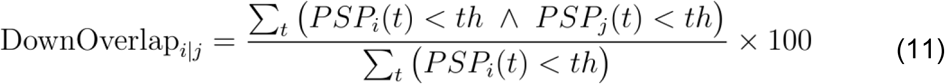

This measure quantifies the fraction of time points where both node *i* and node *j* are in the Down state, relative to the total *Down state* time of node *i*. Thus, the DownOverlap is directional: DownOverlap_i∣j_ is computed with the denominator depending only on node *i*, while DownOverlap_j∣i_ uses the same numerator but normalizes by the *Down states* of node *j*. The DownOverlap reflects the coherence of the SWs processes between the two nodes, with higher values indicating stronger synchronization of their Down *state* transitions.

The Neighborhood %Down (NBR %Down) was calculated in each node to quantify the amount of SW activity arriving from afferent nodes in a given neighborhood of interest. For a given node *i*, it was computed as the average time spent in down states (%Down, eq. 10) of the nodes directly connected to *i*:

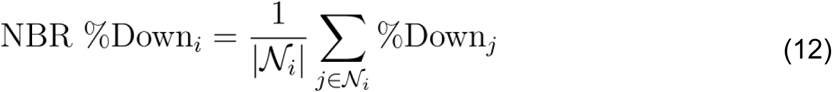

where 𝒩*_i_* denotes a neighborhood of interest for node *i* (specified differently for the toy network and for the whole brain, see below), that is, a set of nodes projecting to node *i*; |𝒩*_i_*| denotes its cardinality (i.e., the number of such nodes).

The *Neighborhood* DownOverlap (NBR DownOverlap) was computed in each node to quantify the temporal overlap of SW activity arriving from afferent nodes in the given neighborhood of interest (see below), thus reflecting the coherence of the SWs processes impinging on a node. For a given node *i*, it was calculated as the average DownOverlap across all ordered node pairs in the given neighborhood of interest 𝒩*_i_* , excluding self-connections:

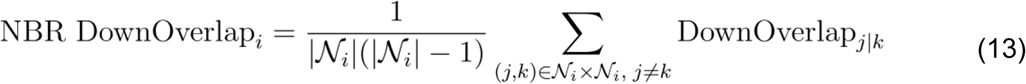

where DownOverlap_j∣k_ denotes the DownOverlap of node *j* relative to node *k*, computed over all ordered pairs of afferent nodes to *i*.

The neighborhood of interest differed for the toy network and for the whole brain model. In the toy network simulations, the neighborhood 𝒩 of the nodes in the distant network was defined by including only afferent connections originating from the perilesional network, thereby focusing only on the impact of perilesional activity on the distant network.

For the whole-brain model, the 𝒩 was computed by thresholding the structural connectivity matrix and retaining only the top 5% of the strongest connections across the network (while the simulation itself used the full, unthresholded connectome). This threshold highlighted a connected subset of the network that included all network nodes, and was chosen to focus on the most influential connections, given the high number of weak connections in the structural connectivity that are unlikely to strongly affect the dynamics. Control analyses varying this threshold showed that results remained stable when retaining between the top 25% and top 1% of the strongest connections.

### 2.7 Structural connectivity metrics

Structural connectivity metrics were used to quantify the effects of the lesion on the spared nodes. The distance from the lesion was computed as the Euclidean distance between each node and the lesion node(s), in parallel to the study of Russo and collaborators^35^. The position of each node is represented in 3D space, with coordinates corresponding to the center of the AAL-defined parcel to which it belongs. The connection loss metric was used to quantify the amount of disconnection that each node suffered following the lesion. It is defined as the percentage of the incoming weights that is lost post-lesion relative to the pre-lesion network. This metric reflects the reduction in excitatory coupling and serves to characterize the degree of disfacilitation caused by the lesion.

### 2.8 Spectral analysis

The power spectral density (psd) of the PSP was computed using the Welch method with time segments of 3 seconds and 50% overlap. From the resulting psd, *δ power* was extracted by summing the power values of frequencies below 4 Hz. To assess the relative change in delta (δ) band between conditions, the *δ ratio* was calculated by dividing the *δ power* in the condition of interest by the *δ power* in the control condition. This ratio provides a normalized measure of the shift in SW activity after the lesion.

### 2.9 Statistical analysis

The bar graphs in this study display the mean values of the reported variables across multiple simulation runs, with error bars representing the corresponding standard deviation.

Exponential fit was performed to model the relationship between *distance from the lesion* and the %Down variable following Russo *et al.*, 2021. The optimal parameters for the exponential decay function were obtained using the *curve_fit* function from the SciPy Python library ^57^. To stabilize variance and enable better linear correlation analysis, the %Down variable was square root–transformed (%Down_sqrt_). Pearson’s correlation coefficient (*r*) was then computed to assess linear relationships between %Down_sqrt_ and each independent variable separately: connection loss, NBR %Down, and NBR DownOverlap.

Partial Least Squares (PLS) regression ^58,59^ was used to combine the independent variables (connection loss, NBR %Down, NBR DownOverlap) into the first PLS latent component (PLS_1_) to predict %Down_sqrt_. This predictive model effectively reduces the dimensionality of the three considered predictors into a single component that is the most linearly related to the outcome variable, and has been used elsewhere in the study of neurophysiology to deal with correlated variables (Colombo et al., 2023). PLS regression was implemented using the *PLSRegression* function from the scikit-learn Python library ^60^. The R^2^ value was computed for both the full model and under 10-fold cross-validation to evaluate generalization performance. The relative importance and contribution of each independent variable to the prediction of %Down_sqrt_ were quantified using the Variable Importance in Projection (VIP) scores for the PLS_1_ component. Variables with VIP scores > 1 are generally considered significant contributors^61^, and the value of 0.8 is often used as a limit below which variables are considered unimportant^58^. Finally, to assess distance-dependent contributions, nodes were divided into quartiles based on their distance from the lesion, and PLS regression was applied separately to each group.

## Results

In this work, we explored two mechanisms proposed to modulate the generation of SWs, namely activity-dependent adaptation and structural disfacilitation. In the context of brain lesions, these mechanisms can be interpreted as proxies for vertical disconnection (e.g., disruption of the ascending activation fibers) and lateral disconnection (e.g., impaired cortico-cortical connectivity). We began by describing how adaptation shapes the dynamics of an isolated cortical population, then examined how SWs emerge through the interaction between adaptation and disfacilitation in coupled populations. Building on this mechanistic foundation, we investigated how lesion-induced disfacilitation and, consequently, functional network-mediated effects drive SW propagation in simplified network motifs. Finally, in a whole-brain model, we quantified the relative contribution of disfacilitation, network hierarchy, and temporal coherence to post-lesional SWs dynamics.

### 3.1 Adaptation induces Up and Down dynamics in an isolated model of the cortical population

The JR model has been extensively used to simulate spontaneous and evoked cortical activity at a mesoscale level (EEG and LFP-like signals)^42,43^. However, it has mostly been applied without considering activity-dependent adaptation dynamics^50^. Here, we characterized the JR model in the presence of adaptation and investigated its relationship with Up and Down dynamics. As a function of the adaptation strength (C), the model qualitatively reproduced distinct dynamical regimes ranging from wake-like states (Up or active state) to those observed during SWS (i.e., slow intermittent Up and Down dynamics). Specifically, for low values of adaptation strength (g=5; Fig. 2A, top-trace), the average postsynaptic membrane potential (PSP) was characterized by an Up (active) state with oscillatory activity at ∼10Hz with an amplitude of ∼7mV, resembling wake-like cortical dynamics^63–65^. By increasing g (Fig. 2A middle traces), we observed a transition to a bistable regime of slow oscillations (SO), where periods of Up states were interspersed by Down states (PSP ∼4mV) together with an increment in the δ power (see psd in Fig. 2A), a fingerprint of SWS dynamics^66,67^. As g further increased, Down states became more frequent and regular^68,69^. At sufficiently high g values, Down states dominated the dynamics over the regular presence of Up states (g=50, Fig. 2A, bottom trace). Notice that a persistent Up state (high-voltage; top trace in Fig. 2A) led to rapid oscillatory patterns, in sharp contrast with a persistent Down state (low-voltage; bottom trace in Fig. 2A).

**Figure 2:**
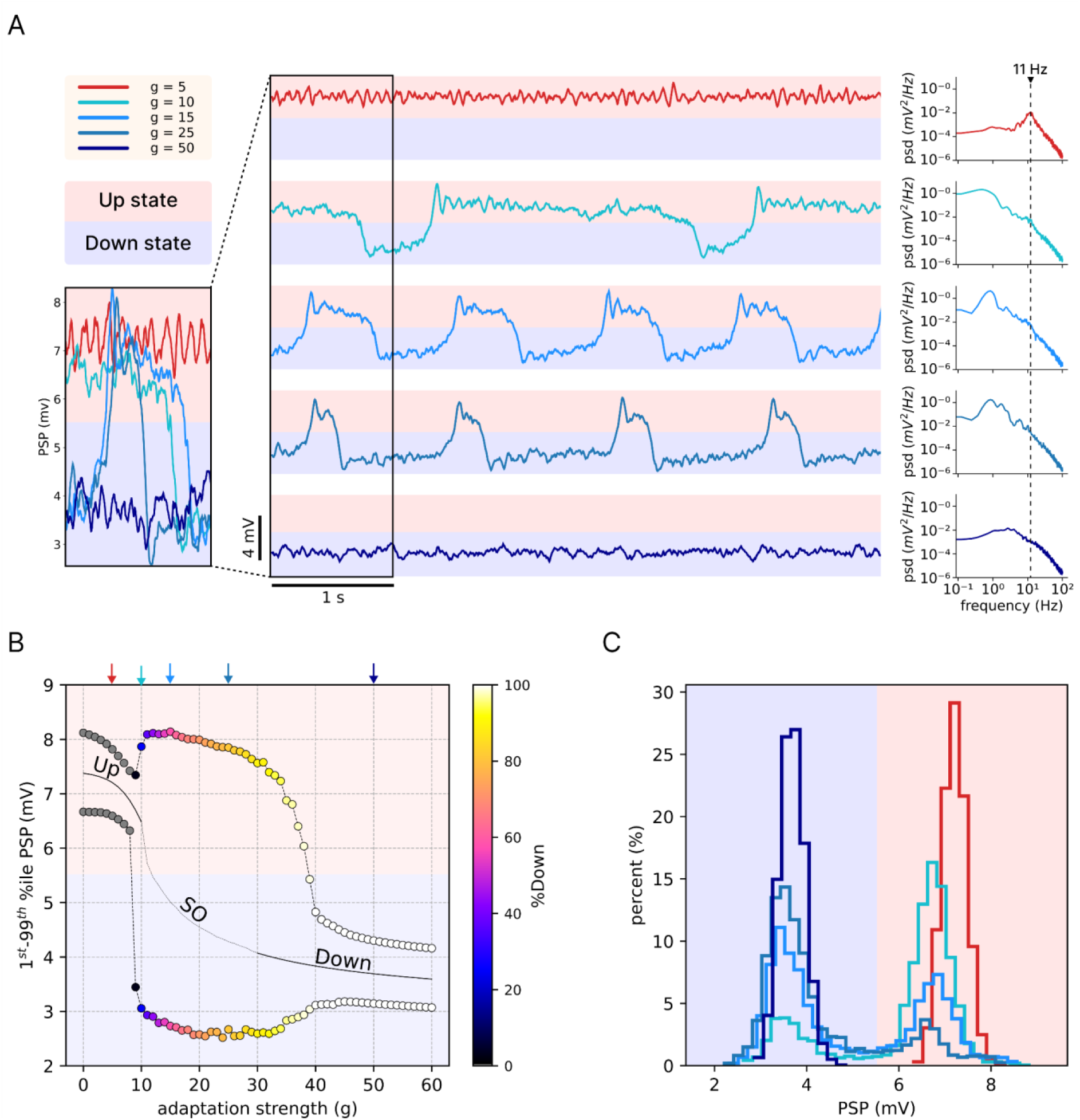
Adaptation induces Up and Down dynamics in an isolated cortical population model. (A) Postsynaptic potential (PSP) traces for different adaptation strength values (*g*). Low *g* (top) leads to persistent Up state, while increasing g induces slow oscillations (SO), characterized by alternating Up and Down states. For high *g* (bottom), the system remains in a persistent Down state. Right panels show the power spectral density (psd) for each case. (B) Bifurcation diagram showing the 1st and 99th percentiles of PSP as a function of *g*. Three distinct dynamical regimes are observed: persistent Up state (gray dots, left), SO (colored region, middle), and persistent Down state (white dots, right). The color scale indicates the percentage of time spent in the Down state (%Down). The solid black line indicates the presence of a stable fixed-point attractor, corresponding to a high-voltage Up state for low values of *g* (left) and a low-voltage Down state for sufficiently high *g* (right). The vertical arrows above the plot denote the cases illustrated in panel (A), with colors matching the corresponding adaptation strength *g*. (C) Distribution of PSP values for the five cases shown in (A). At low and high g, the distribution is unimodal, reflecting stable Up and Down states, respectively. At intermediate g, a bimodal distribution emerges, corresponding to SO dynamics.

Then, we explored the model’s dynamical landscape by systematically varying g from 0 to 60 and quantified the amplitude of oscillations as 1st and 99th percentiles of the PSP (Fig. 2B). The Up state (high-voltage/active state) regime corresponded to a stable fixed-point (Fig. 2B solid line), which, when subjected to stochastic noise, exhibited low-amplitude alpha oscillations (from ∼6.5 mV to ∼8.2mV; Fig. 2B gray dots; Fig. S1). For instance, when *g*=0, the system was in a subcritical regime near a supercritical Hopf bifurcation (Fig. S1) and acted as a damped oscillator, decaying towards the fixed-point attractor. This behavior persisted for small increases in g (g<9; Fig. 2 and S1). However, for larger values of g (g>9 up to 39), the model exhibited large oscillations between high- and low-voltage states, driven by the slow negative feedback effect of adaptation (Fig. 1C). As firing activity continues, the adaptation gradually builds up, lowering sensitivity to input potentials, until the Up state loses stability, triggering a transition to the Down state. In this low-activity regime, adaptation slowly decays until it reaches its minimum, enhancing sensitivity to input potentials, allowing the system to spontaneously generate a new Up state, thus re-initiating the Up/Down cycle. For high values of g (>39), the system fell in a persistent Down state, due to a stable low-voltage fixed point attractor (Fig. 2B and S1B). Notably, near the transition, from a persistent Up to SO (g∼10) or from SO to persistent Down (g∼40), the system remains at a stable-fixed point (solid line in Fig. 2B), however, due to the stochastic noise, transitions were still observed (Fig. S1B).

**Figure 3.**
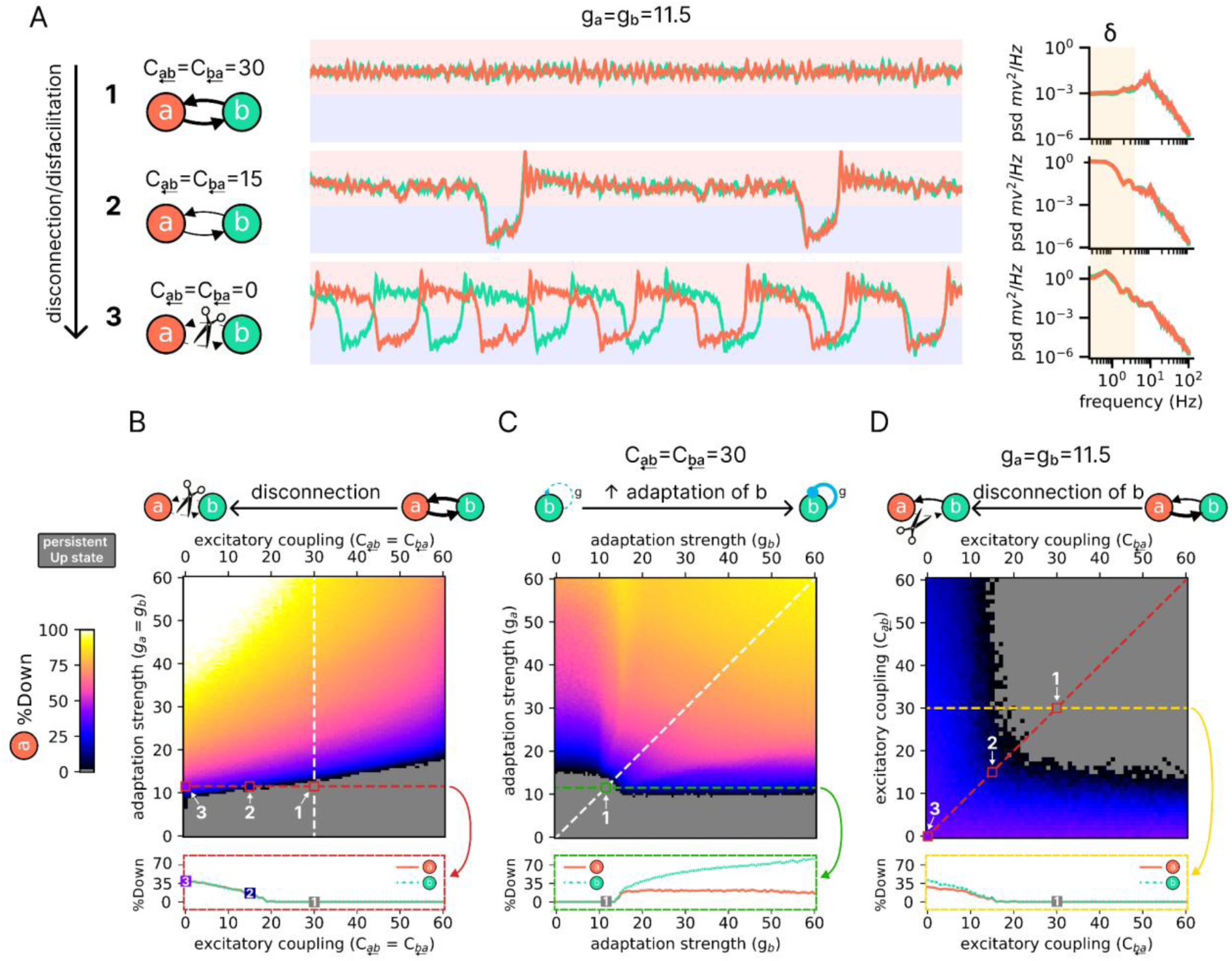
Effects of adaptation strength and excitatory coupling on slow wave emergence in two coupled populations. (A) Example dynamics of two symmetrically coupled populations (JR_a_ in orange, JR_b_ in cyan) for three levels of excitatory coupling (C_ab_=C_ba_). With C=30 and g=11.5 (top, #1), both populations show alpha oscillations in a persistent Up state. Reducing coupling to C=15 (middle, #2) induces a transition to slow oscillations (SO), reflected in an increase in δ power. Complete disconnection (C=0, bottom, #3) further enhances SO activity. Right panels show the corresponding power spectral densities (psd), with the δ band highlighted by the yellow span. (B) Heatmap showing the percentage of time spent in the Down state (%Down) in JR_a_ as a function of adaptation strength (g, y-axis) and symmetric excitatory coupling (C, x-axis). The red horizontal dashed line highlights the effect of reducing C at g=11.5, revealing a transition from a persistent Up state (gray region) to SO (colored region). Bottom panel: %Down along the red horizontal dashed line, illustrating the progressive reduction of SO with increasing C. (C) Effect of asymmetric adaptation. Heatmap shows %Down in JR_a_ as a function of g_b_ (x-axis) and g_a_ (y-axis), with symmetric coupling fixed at C=30. The white diagonal dashed line corresponds to the vertical white dashed line in panel A. The green horizontal dashed line indicates that varying g_b_ alone can induce a transition from a persistent Up state to SO in JR_a_, despite g_a_ being constant. Bottom panel: %Down along the green dashed line, showing that in JR_b_ (where g varies) the time in the Down state increases continuously, whereas in JR_a_ (where g is fixed) it increases but quickly reaches a plateau. (D) Effect of asymmetric coupling. Heatmap shows %Down in JR_a_ as a function of C_ba_ (x-axis) and C_ab_ (y-axis), with g fixed at 11.5. The diagonal red dashed line corresponds to the horizontal red dashed line in panel B. The yellow dashed line illustrates that reducing C_ba_ (i.e., disconnecting JR_b_) alone is sufficient to induce a transition from a persistent Up state to SO in JR_a_, even when C_ab_ remains constant. Bottom panel: %Down along the yellow dashed line, emphasizing the dependence of the transition on directional disconnection. Numbers 1, 2, and 3 in all panels correspond to identical parameter configurations across plots. The corresponding heatmaps for population JR_b_ is shown in Supplementary Fig. S2.

Finally, in Fig. 2C, we report the PSP for the five cases illustrated in Fig. 2A. At the two extremes (g=5 and g=50), the PSP exhibited a unimodal distribution, corresponding to the high-voltage attractor of the Up state and the low-voltage attractor of the Down state, respectively (Fig. 2C). Conversely, for intermediate values of g, the PSP followed a bimodal distribution, reflecting the alternation of Up and Down states typical of SO. Altogether, our results demonstrate that, in a local circuit (Fig. 1D I), represented by the JR model, manipulating the strength of adaptation induces a phase transition from a persistent Up state to a bistable SO regime, resembling a transition from wakefulness to sleep-like states.

### 3.2 Effects of adaptation strength and excitatory coupling on slow oscillations in two connected populations

The cerebral cortex relies on recurrent excitatory connectivity to sustain stable activity, and its disruption profoundly alters neuronal dynamics. Consistent with empirical evidence showing SO after cortical deafferentation and brain injury^35,70–72,11^, we hypothesized that in our model sleep-like dynamics may arise from two complementary mechanisms: (i) *structural disfacilitation*, reflecting reduced cortico-cortical afferent excitatory input^31,32,34^, and (ii) *loss of ascending neuromodulatory drive*, which reduces acetylcholine and norepinephrine release, thereby enhancing neuronal adaptation and favoring SO emergence^73^. To test this, we coupled two JR neural populations (JR_a_, JR_b_) through bidirectional excitatory connections without delays (Fig. 1D II) and systematically varied the adaptation strength (g) and excitatory coupling (C) (Fig. 3).

We first examined the dynamics of two symmetrically coupled populations, JR_a_ and JR_b_, with excitatory coupling strength fixed at 30 (C_ab_=C_ba_=30) and g=11.5. Under these conditions, both populations exhibited alpha oscillations in a persistent Up state (Fig. 3A, top trace and PSD; #1). Reducing C induced a transition from the persistent Up state to SO, characterized by an increase in δ power (Fig. 3A, middle trace; #2). This effect can be interpreted as an increase in the “level of structural disfacilitation”^74,75^. When these populations were fully disconnected (C=0), SO activity became more frequent and regular (Fig. 3A, bottom trace; #3). We then systematically investigated the impact of disconnection across different levels of adaptation strength (Fig. 3B). As highlighted by the red horizontal dashed line, decreasing C (x-axis) while keeping g (y-axis) fixed revealed a phase transition from SO (colored region) to a persistent Up state (gray region). For example, when uncoupled (C=0), both populations displayed SO, spending approximately 35% of the time in the Down state (%Down) (Fig. 3B, bottom). With increasing coupling strength, the %Down progressively decreased, disappearing around C≈20, beyond which the system settled into a stable persistent Up state (Fig. 3B, bottom).

Next, we explored the case of asymmetrical variations of g, i.e., whether manipulating g in only one population could induce a change in the dynamics of the other population, while keeping C=30 fixed (Fig. 3C). The bifurcation diagram for population JR_a_, as a function of g_a_ and g_b_, revealed that a transition from persistent Up state (gray area) to SO (colored area) could occur by simply manipulating the adaptation in the other population, JR_b_. For instance, when g_a_=11.5 and g_b_ was varied, population JR_a_ transitioned from a persistent Up state to SO (Fig. 3C, green horizontal dashed line). Importantly, the %Down in JR_b_, increased as a function of g_b_ while, in the connected population JR_a_ (where g was fixed), the %Down reached a plateau (Fig. 3C, bottom).

Finally, we evaluated the case of asymmetrical variations of C. We showed that for fixed values of g in both populations JR_a_ and JR_b_ (g_a_=g_b_=11.5), population JR_a_ can transition from a persistent Up state to SO as a function of C_ba_ (the coupling strength from JR_a_ to JR_b_) even if C_ab_ (the coupling strength from JR_b_ to JR_a_) remains unchanged (Fig. 3D). In other words, the disconnection of population JR_b_ induces a transition from a persistent Up state to SO in JR_a_. Moreover, we replicated these results in the Wilson-Cowan rate model, demonstrating that the observed dynamics are consistent across different models (Supplementary Material; Fig. S3).

### 3.3 Slow wave propagation in a toy-network model

In the previous section, we demonstrated that the transition from a persistent Up state to SO depends on local ascending (e.g., adaptation level) and cortico-cortical (e.g., coupling strength) connections in a simple bidirectionally coupled system. To further explore these aspects, we extended the model to include the interaction with other regions (nodes), incorporating a distance-dependent connectivity pattern to mimic both local and long-range connections. This toy network was composed of two sub-networks, referred to as the perilesional network and the distant network, in accordance with where the lesion was applied (Fig. 4A). The control pre-lesion case (Fig. 4B) represents the intact network (i.e., before any disconnection was applied) in the wake-like (active) regime, where all the nodes oscillated at ∼10Hz. In this toy network, adaptation was kept fixed (g=11.5) and the total excitatory input coupling was equalized across nodes (c=30; see Methods 2.4) corresponding to configuration 1 in Fig 3A.

**Figure 4.**
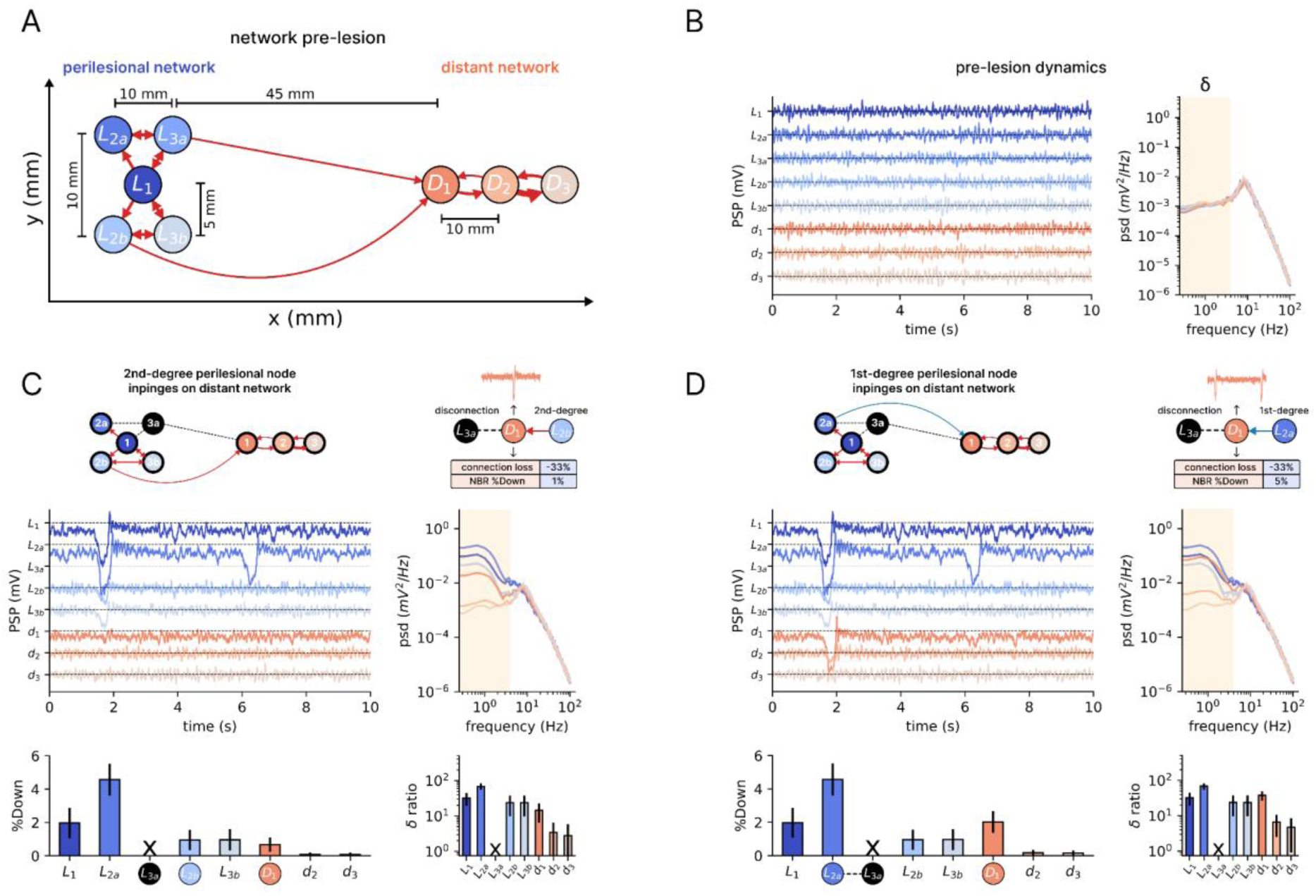
Impact of disconnection and disfacilitation on slow wave (SW) propagation in a toy network. (A) Schematic representation of the toy network model, composed of a perilesional network (blue) and a distant network (red), here shown in the pre-lesion control condition (i.e., before performing any disconnection). Nodes are connected with a distance-dependent connectivity pattern, with the thickness of the arrows indicating coupling strength. The perilesional network projects to the distant network via node D1. Delays between nodes are determined by inter-node distances and a fixed propagation velocity of 4 m/s. (B) Activity and spectrum of the control condition: all nodes oscillate in the alpha band in a persistent Up state (awake-like regime). Left panel shows postsynaptic potential (PSP) traces recorded from perilesional (blue) and distant (red) nodes. The right panel displays the corresponding power spectral density (psd), highlighting the dominant alpha activity. The δ band is highlighted by the yellow span. (C) Effect of disfacilitation following the lesion of node L3a (black node), which projected both locally and to the distant network in the control condition. The loss of L3a primarily affects its first-degree nodes L1 and L2a, with L2a exhibiting the strongest impact due to additional inputs from its first-degree node L1. The distant node D1 also shows signs of SW propagation, since now it receives inputs from node L2B (a second-degree node relative to the lesioned node L3a). The bottom panels quantify the percentage of time spent in the Down state (%Down) and the δ ratio for each node. The δ ratio was calculated by dividing the δ power in the network after the lesion by the δ power in the control condition. The percentage of time neighboring nodes (presynaptic nodes) spend in Down states (here, 1% for D1) is referred to as NBR %Down (see Methods 2.6). (D) Effect of modifying connectivity so that D1 receives input from node L2a rather than node L2b, thus from a first-degree node relative to the lesioned node (NBR %Down=5%) instead of a second-degree node, as in (C) (NBR %Down=1%). This change amplifies SW propagation, leading to a greater impact on distant nodes (D2 and D3), as reflected by increased time spent in the Down state and a higher δ ratio.

Within this framework, we explored the impact of disfacilitation over the perilesional network and its influence on the distant network, by simulating a virtual local lesion, removing a node and all of its connections. Specifically, the lesion was simulated by disconnecting a node (L3a) that projected to both local nodes (i.e., perilesional network) and distant nodes (i.e., distant network) (see Fig. 4C). The impact of disfacilitation was quantified by the amount of time spent in the Down state and by the δ ratio (Fig. 4C-D).

Within the perilesional network, the nodes most affected were those directly connected to the lesioned node (L2a and L1, hereafter referred to as first-degree neighbors), though to different extents. Node L2a was the most impacted, as it not only lost its direct connection to the lesioned node but also received input from L1, which itself was directly connected to the lesion (i.e., a first-degree neighbor; see Fig. 3D). In contrast, node L1 was less affected, suggesting that its stability depended less on the lesioned node. This reduced impact may be explained by the small recurrent circuit involving L1, L2b, and L3b, which received only indirect (second-degree) input from the lesion.

Next, we examined the impact of disfacilitation on the distant network. The perilesional network projects only to node D1 in the distant network (Fig. 4A). In the paradigm depicted in Fig. 4C, D1 lost its connection to the lesioned node and received inputs from node L2b, a second-degree neighbor of the lesion (less affected). To characterize these two aspects, we computed the connection loss, which represents the proportion of lost connections (coupling strength) relative to the control condition for each node (-33% for D1), and the NBR %Down, which quantifies the time neighboring nodes spend in Down states (1% for D1) indicating the extent to which inputs come from affected regions. Although less pronounced, the effect on the D1 dynamics resembled that observed in L1, characterized by a small amount of time spent in the Down state and increased δ ratio. Interestingly, we observed signs of SW propagation in the distant network, as indicated by time spent in the Down state and δ ratio of D2 and D3, which received progressively indirect inputs from the lesion, specifically the second- and third-degree neighbors of the lesion, respectively (Fig. 4C).

Finally, we examined the effect on D1 when it received input from a first-degree neighbor of the lesion (NBR %Down=5%, Fig 4D), rather than from a second-degree neighbor of the lesioned node (as in Fig 4C). As shown in Fig. 4D, this configuration resulted in a greater impact on the distant nodes and enhanced SW propagation to nodes D2 and D3. These findings suggest that the spread of SWs to distant nodes depends not only on the presence of SWs in the perilesional network but also on the hierarchical position of the affected nodes relative to the lesion, which is mirrored in their level of SW activity or δ power.

### 3.4 Slow wave coherence drives propagation in a toy-network model

To further investigate the impact of lesion-induced SWs, we examined how their temporal organization influences propagation to the distant network. Specifically, we compared two conditions following the lesion of the L1 node: two uncorrelated SW-generating nodes (Fig. 5A) and two correlated SW-generating nodes (Fig. 5B). Even though the amount of SW activity conveyed to the D1 node was identical between conditions (NBR %Down=14%), the coherence of SW activity across nodes led to remarkably different effects. When SW activity was coherent, the distant nodes (D1, D2, D3) were significantly more impacted, as reflected in the increased time spent in the Down state (%Down) and a higher δ ratio. These observations suggest that the propagation of SW activity into intact nodes depends not only on the amount of SWs (NBR %Down) but also on the coherence of these oscillations, highlighting a dependence on the temporal dynamics of the perilesional network.

To quantify the impact of coherent SWs on distant nodes, we computed the percentage of Down state overlap (DownOverlap) between pairs of perilesional nodes (see Methods 2.6; Fig. 5C,D). Focusing on the coherence among nodes projecting to D1, we first calculated the DownOverlap between neighboring pairs (L2b–L3a and L2a–L3a), corresponding to the two scenarios illustrated in Fig. 5A,B. We then averaged the DownOverlap across all neighbors of D1 to obtain the NBR DownOverlap measure. As expected, L2a and L3a exhibited markedly higher Down state overlap (NBR DownOverlap = 94.2% ± 1.2%) than L2b and L3b (NBR DownOverlap = 11.7% ± 4.6%; Fig. 5E), resulting in a stronger impact on D1 and consequently on D2 and D3.

Finally, to assess the causal role of SW activity coherence in modulating SW propagation in the distant network, we altered the propagation delay from L2a to D1 (as in the scenario depicted in Fig. 5B), thereby reducing the overlap of the two SW inputs arriving at D1 (see Fig. 5F). As the delay increased (less overlapping of the SW activity impinging to D1), we observed a progressive decrease in the time spent in the Down states in the distant network, suggesting that SW coherence plays a key role in network-wide SW activity propagation.

**Figure 5.**
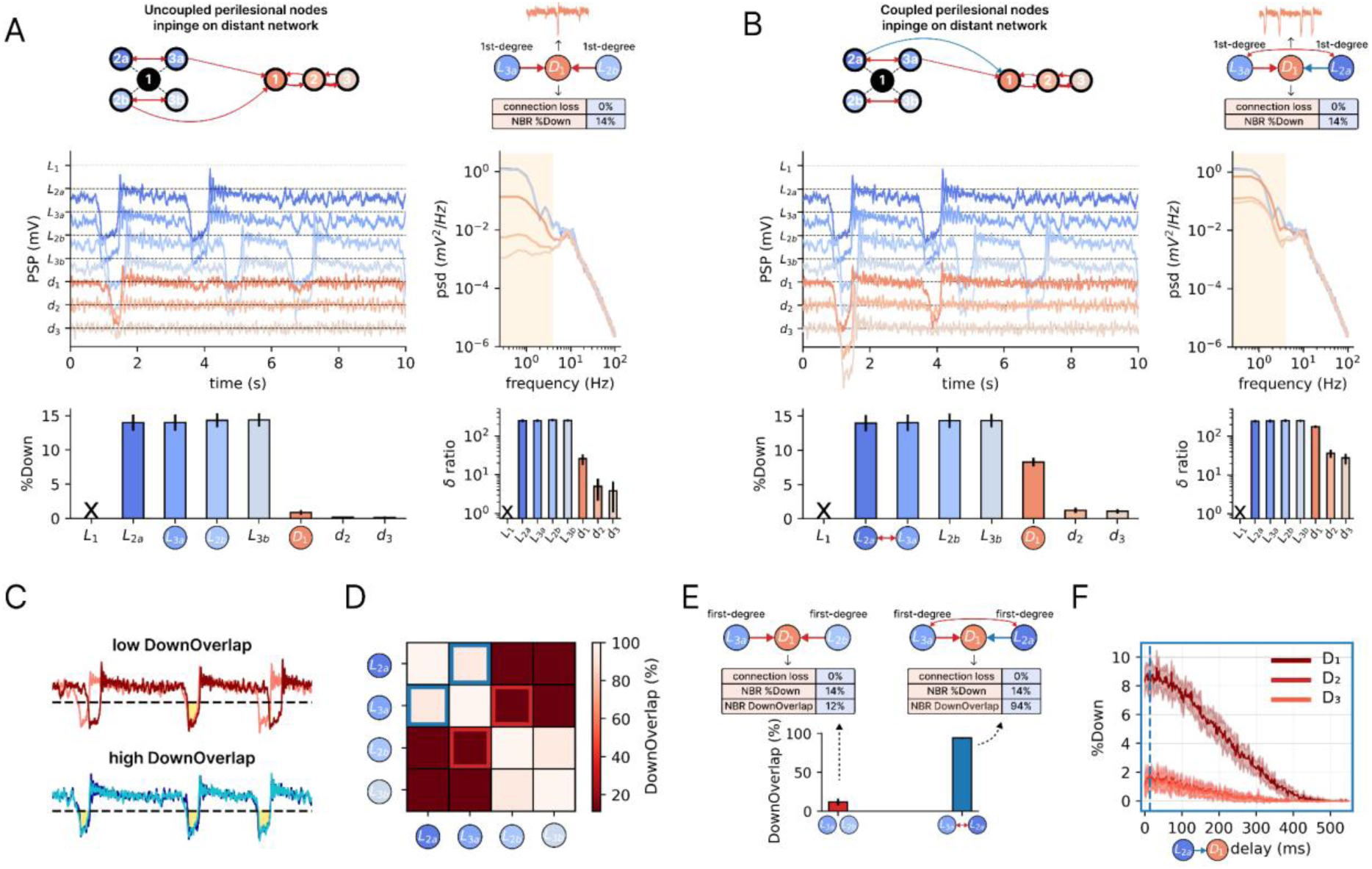
Impact of coherent and uncoherent slow wave inputs on the propagation of activity in a distant network. (A) Schematic representation of the network model where uncoupled perilesional nodes impinge on the distant network, resulting in uncoherent SW inputs. Below, postsynaptic potential (PSP) traces of the network, with the perilesional nodes (blue) and distant nodes (red). Right, corresponding power spectral density (psd) and δ ratio. The δ band is highlighted by the yellow span in the psd. (B) Same as in (A), but the perilesional nodes that impinge on the distant network are coupled, resulting in coherent SW inputs. For the same amount of SW input (i.e., same NBR %Down for D1 in both scenarios), coherent inputs lead to a greater impact on the distant network, as reflected by the increased time spent in the Down state (%Down) and higher δratio. The δ ratio was calculated by dividing the δ power in the network after the lesion by the δ power in the control condition. (C) Illustration of DownOverlap calculation. Given a pair of PSP traces, the *DownOverlap* computes for each trace the fraction of its total Down state duration during which both traces simultaneously fall below the Down state threshold (black dashed horizontal line). (D) Heatmap representing the *DownOverlap* between pairs of perilesional nodes. Lighter colors indicate higher overlap, showing that specific node pairs exhibit stronger coherence (blue and red squares). (E) The L2a–L3a node pair (corresponding to the coupled perilesional nodes scenario in B) shows significantly higher overlap than the L2b–L3a node pair (the uncoupled perilesional nodes scenario in A). The average DownOverlap of presynaptic nodes projecting to a given postsynaptic node (here, D1) is referred to as NBR DownOverlap (see Methods 2.6). Given that DownOverlap is not necessarily symmetric (i.e., the overlap from node *i* to node *j* may differ from *j* to *i*), we reported the average of the two directions. (F) Effect of propagation delay on Down state occurrence in the distant network. Delay was progressively increased between L2a and D1 (the coupled perilesional nodes scenario in B ), reducing the overlap of the two SW activity inputs arriving at D1. Traces represent the mean over 10 runs, and the shaded area indicates the standard deviation. The blue vertical line marks the actual delay in (B).

### 3.5 Slow wave propagation in a whole-brain network model

So far, we have explored the emergence of SWs and their propagation to other nodes in a toy network. In doing that, we identified three key factors: i) the role of disconnection (disfacilitation) in SW emergence, ii) the dependence on hierarchical node position relative to the lesion, which directly impacts the amount of SW activity (or δ power) of a given node, and iii) the coherence of SW activity from different affected nodes impinging on the intact nodes, i.e., the dependence on their temporal profile. To examine these dynamics at a larger and realistic scale, we extended our analysis to a whole-brain model.

We implemented the JR model in a whole-brain composed of 80 nodes^54^ (Fig. 6). Similar to our control toy network, the whole-brain network dynamics in the control condition were characterized by alpha oscillations (Fig. 6A, gray traces). To assess the impact of lesions, we systematically disconnected each node and examined its effects on both perilesional and distant nodes. As shown in Fig. 6A for a representative lesion (black dot), the most affected nodes were the perilesional ones, where Down states emerged along with an increase in δ power (Fig. 6A). Moreover, we showed that SW propagation decayed exponentially with the distance from the lesion, i.e., closer nodes were more affected than distant ones, consistent with empirical observations (Fig. 6B)^35^.

**Figure 6.**
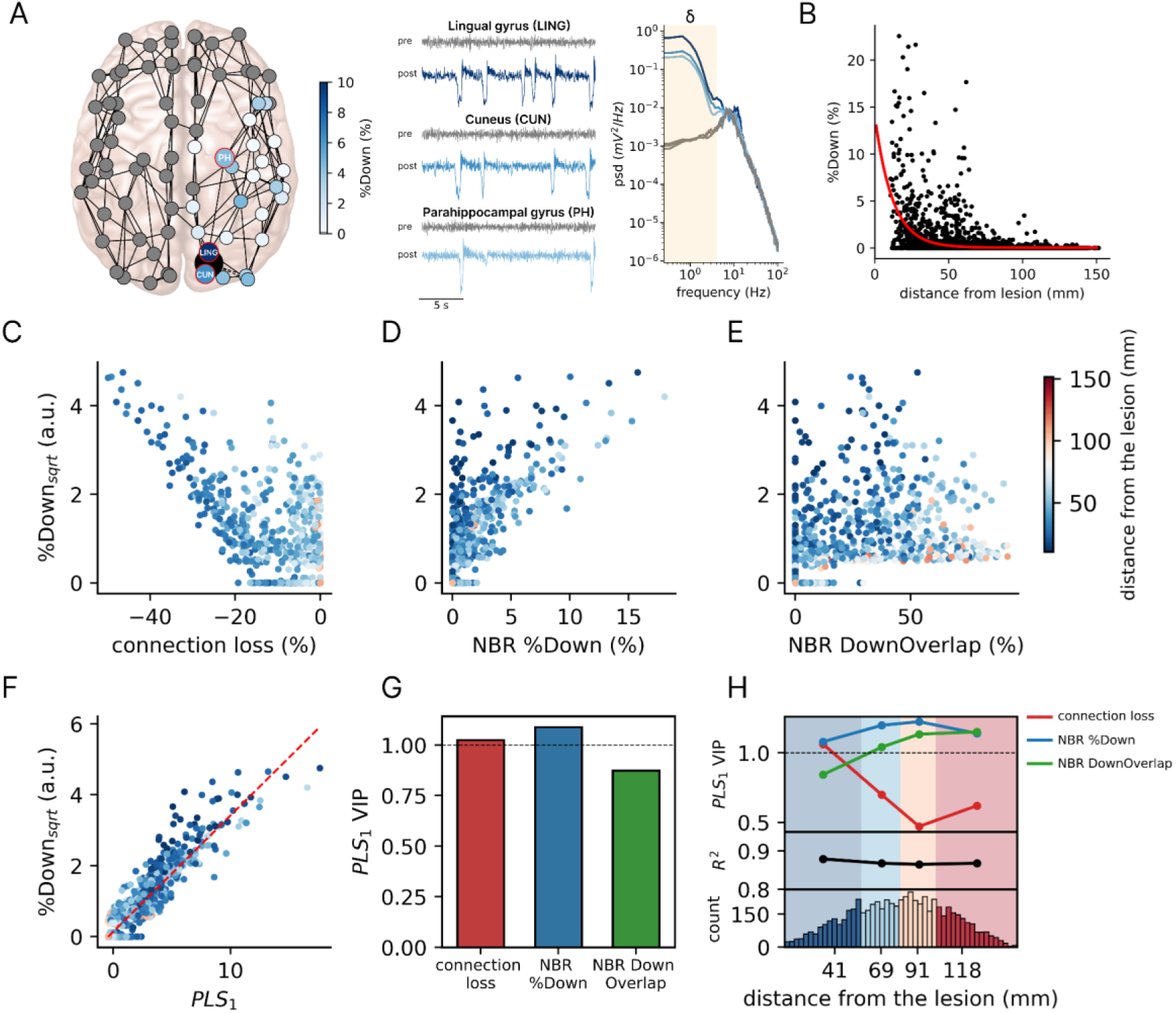
Whole-brain modeling of slow wave generation and propagation following lesion. (A) Whole-brain network model with nodes defined by the Automatic Anatomical Labeling (AAL) atlas ^55^. The lesion of a node (here, the right calcarine cortex) is represented as a black dot, with the impact on the spared nodes quantified as the time spent in the Down state (%Down). The node colors indicate %Down, as shown in the accompanying color bar, with gray nodes representing regions where no Down states were detected. Traces illustrate simulated PSP activity from three representative regions in the pre-lesion (control; gray) and post-lesion (blue hues) conditions. In the control condition, the nodes oscillate in the alpha band in a persistent Up state (awake-like regime). After the lesion, Down states emerge along with an increase in δ power in power spectral density (psd). The δ band is highlighted by the yellow span in the psd. (B) Percentage of time spent in the Down state (%Down) as a function of the Euclidean distance from the lesion. The red curve represents an exponential fit (R^2^=0.21), indicating a distance-dependent decay in SW propagation, consistent with experimental results ^35^. (C) Relationship between connection loss (percentage of lost coupling strength due to the lesion relative to control) and the square root of %Down (%Down_sqrt_). A negative relationship is observed (r=-0.76). (D) Relationship between the time spent in the Down state of neighboring nodes (NBR %Down) and %Down_sqrt_. A positive relationship is observed (r=0.81). (E) Relationship between the DownOverlap of neighboring nodes (NBR DownOverlap) and %Down_sqrt_. The variables show a positive relationship (r=0.67). (F) Partial least squares regression (PLS) of %Down_sqrt_ and the score of the first PLS component (PLS_1_) derived from the three independent variables (connection loss, NBR %Down, and NBR DownOverlap). The model explained ∼90% of the variance (R^2^=0.89; after cross-validation R^2^=0.88). (G) PLS1 weights for the three independent variables, indicating their relative contributions. (H) Distance-dependent contributions of the three independent variables to %Down_sqrt_. Nodes were divided into quartiles based on their distance from the lesion, and PLS regression was performed for each subset. For each quartile, the model’s R^2^ and the VIP scores of each variable are reported.

To further explore the impact of disconnection following the lesion, we computed, for each node, the fraction of connection loss, as done in the toy network. We observed that the square root of the time spent in the Down state (%Down_sqrt_; see Methods 2.9) followed a negative relationship with the fraction of connection loss, i.e., nodes more connected to the lesion site were impacted the most (Fig. 6C). Next, we assessed the dependence of SW propagation on the hierarchical node position relative to the lesion. Specifically, we explored whether the time spent in the Down states could be explained by the time spent in the Down states by neighboring nodes (NBR %Down). We observed a positive relationship, suggesting that recurrent connectivity, i.e., the influence of neighboring nodes, can explain, to some extent, the impact on a given node (Fig. 6D). Finally, we assessed whether the amount of SW overlap in presynaptic nodes could account for the time spent in the Down state of the postsynaptic nodes (NBR DownOverlap). Albeit more dispersed, we also observed a positive relationship between the amount of SW overlap and the %Down_sqrt_ (Fig. 6E).

However, it is important to note that, while each of the three explored predictors (connection loss, NBR %Down, and NBR *DownOverlap*) were significantly correlated with %Down_sqrt_ (Fig. 6C-E), each predictor exhibited a clear dispersion around zero on the x-axis, suggesting that neither predictor alone fully accounts for the variability in the time spent in the Down states. To further investigate the combined contribution of these predictor variables in explaining the time spent in the Down state and SW activity propagation, and also to address potential multicollinearity, we employed partial least squares regression (PLS). Specifically, we aggregated the three factors into the first PLS latent component (PLS_1_) and subsequently related its score to %Down_sqrt_. We observed a strong linear relationship between the PLS_1_ and %Down_sqrt_ (Fig. 6F), with all factors robustly explained the variance of the outcome, as indicated by their VIP scores (Fig. 6G; all variables have scores around 1; see Methods 2.9). Finally, we found that these predictors exert distance-dependent effects: connection loss had higher VIP scores for nodes closer to the lesion (stronger predictive power for %Down_sqrt_), NBR DownOverlap had higher VIP scores for more distant nodes, and NBR %Down showed VIP scores largely independent of distance from the lesion (Fig. 6H). These findings were also consistent for a different model (Supplementary Material; Fig. S4).

## 3. Discussion

In this study, we employed a population model of cortical activity and applied a multiscale analytical framework to examine the mechanisms underlying the emergence and propagation of SWs following brain lesions (Fig. 1). Our findings identify three interacting factors: i) the critical role of disconnection, leading to increased adaptation and/or disfacilitation, in SW generation, ii) the hierarchical organization of cortical connectivity as a constraint on SW propagation, and iii) temporal coherence as a recruitment modulator of distant structurally intact nodes. Together, these results reveal that post-lesional SW propagation depends jointly on topological distance from the lesion and dynamic synchrony of SWs, underscoring the interplay between local connectivity disruption and global network organization in shaping cortical dynamics.

SWs are a hallmark of cortical dynamics during NREM sleep. Organized into a slow, quasi-periodic rhythm, they reflect the alternation between periods of neuronal firing (Up states) and periods of hyperpolarization associated with neuronal silence (Down states, or off-periods). During NREM sleep, SWs and the associated off-periods disrupt network interactions leading to loss of consciousness, while they simultaneously promote restorative and homeostatic processes^76–80^. When SWs intrude into physiological wakefulness, they can impair motor and cognitive functions in a topology-specific manner, as shown in sleep-deprivation studies^21,23,24^. Crucially, SWs have also been observed during wakefulness following brain lesions of varying extent and etiology^10,81^, where they can produce substantial network and functional impairments^72,82–84^. Hence, characterizing the mechanisms responsible for the generation and propagation of post-lesional SWs are essential for understanding the functional consequences of brain injury and for identifying new therapeutic targets.

To address this issue, we systematically explored the effects of virtual lesions and structural disconnections on neuronal dynamics. Specifically, we considered that lesions can disrupt both ascending fibers, leading to increased adaptation (g), and cortico-cortical fibres (C), leading to disfacilitation. We thus varied parametrically and regionally both adaptation and excitation and tested the effects of these local manipulations on neuronal activity using a multiscale approach, from an isolated population up to a human whole-brain connectome (Fig. 1).

We first probed the dependence of the model on global changes in adaptation and disfacilitation in a simple single- and two-populations model. As expected, uniformly increasing adaptation resulted in the cyclic, system-wide alternation between Up and Down states across the whole system (Fig. 2 and Fig. 3B). Comparable effects were observed following symmetrical (>50%) removal of cortico-cortical excitatory interactions (Fig. 3A-B). The effects of these global manipulations across nodes generalized previous single-node bifurcation diagrams^69^ and recapitulated the main electrophysiological features found in N3 sleep, as well in conditions of massive deafferentation, such as in cortical slices^85^, slabs^70^, hemispherotomy^86^ and severe brain injury^87,88^.

We next turned to our primary aim: quantifying how local lesions and discrete disconnections shape network-level neuronal activity. We first modeled a local disruption of ascending fibres by increasing adaptation at a single node within a small network. We found that once adaptation crossed a critical threshold, functional effects extended beyond the structurally deafferented node: recurrent silent periods projected onto the distant node shifted its bifurcation landscape promoting a transition to SO regimes (Fig. 3C). We then explored the effects of the selective removal of cortico-cortical afferent connections. Reducing excitatory contacts to the target node shifted its activity toward the SO. Also in this case, this local transition produced effects on the activity profile of the distant node (Fig. 3D). Thus, local manipulations in a two-population model point to two complementary routes to post-lesional SWs: (i) a primary structural disconnection of ascending and/or cortico-cortical afferents, that directly increase adaptation and/or disfacilitation, leading to the generation of local SO; (ii) secondary, functional network-mediated effects whereby the silent periods generated by the primary disconnection reshape the dynamical landscape of distant nodes.

We then examined these structural and functional effects from a topological perspective using more complex networks, incorporating a distance-dependent connectivity pattern among multiple populations to mimic both local and long-range connections (Fig.4A). Here, we focused primarily on the functional impact of cortico-cortical disconnections for two reasons. First, the consequences of disrupting lateral excitation (i.e., disfacilitation) for SWs are less well understood than those of altering ascending neuromodulation^12,70,89^. Second, focal cortical and subcortical lesions are more likely to compromise cortico-cortical pathways, which vastly outnumber ascending projections^90^.

We found that lesioning a single node primarily altered activity in its nearest neighbors—those experiencing the greatest loss of connections and disfacilitation—shifting their dynamics from wake-like activity (Fig. 4B) to a regime dominated by SO (Fig. 4C). Importantly, these changes also propagated to more distant nodes and scaled with hierarchical distance from the lesion. This pattern reflects both primary structural disconnection and secondary functional network effects: nodes already structurally deafferented were more likely to be affected when they also received inputs from nodes with first-degree connections to the lesion. As illustrated by the comparison between Fig. 4C and Fig. 4D, receiving direct input from these sources of prominent SWs increased the likelihood that a distant node would exhibit Down states, even when the degree of structural disconnection was held constant.

Finally, probing lesions at this scale revealed a third mechanism by which neuronal activity is altered beyond directly deafferented nodes. Here, the emergence of SWs in a distant node depended on the synchrony of SO arriving from nodes topologically closer to the lesion. This timing-dependent effect was evident even when the total convergent SW input was held constant. As illustrated by Fig. 5A versus Fig. 5B, distant nodes receiving input from perilesional populations with incoherent SO behaved differently from those driven by coherent perilesional sources shaped by local reciprocal connectivity. In the latter case, coherent projections from a tightly connected perilesional focus were sufficient to promote further percolation of SWs, extending to nodes several steps away from the lesion. Thus, a node’s vulnerability to SW intrusion scales with both the strength and coherence of SW activity in its presynaptic neighborhood.

Overall, analysis of a topologically organized cortical motifs revealed three concurrent mechanisms governing the generation and propagation of post-lesional SWs: first, primary structural disconnection; second, network-mediated effects that scale with the input received from the perilesional SW focus; and third, a timing-dependent effect governed by the synchrony of those inputs. To examine how these mechanisms interact under more realistic conditions, we applied the same manipulations (i.e., node lesions) to a human whole-brain connectome (Fig. 6A). Indeed, in a complex brain network with a hierarchical, recurrent, and modular architecture^91,92^, these mechanisms are likely to be strongly interdependent. For example, nodes disconnected by a lesion are typically embedded within neighborhoods that are likewise compromised, where prominent SWs may be further amplified by synchrony. To disentangle their individual contributions, we applied partial least squares (PLS) regression and derived a latent component capturing all three factors. This composite measure explained ∼90% of the variance in post-lesional SW activity (Fig. 6F). Notably, all three factors contributed to the emergence and propagation of SWs, but their effects were distance-dependent: structural disconnection primarily accounted for prominent SW intrusions in nearby nodes, whereas SW coherence better explained propagation of smaller events to more distant regions (Fig. 6G–H).

The spatial gradient predicted by our computational model not only aligns with classic reports of prominent perilesional EEG slowing^10,26,27^ but also provides a mechanistic account of intracranial recordings of SW dynamics following controlled radiofrequency thermocoagulation (RFTC) lesions^35^. In those recordings, SW intrusion peaked near the lesion, exhibited long-range propagation to connected sites, and decayed exponentially with anatomical distance—consistent with our modeling results (Fig. 6B). This hierarchical gradient mirrors the exponential distance rule of cortical connectivity^93,94^ and likely reflects SW spread via cascades of structural and functional effects, whereby upstream Down states induce downstream neuronal silencing, as suggested by our model.

These modeling results position SWs as the potential electrophysiological substrate of diaschisis, the classical concept introduced by von Monakow in 1914^8^. Von Monakow originally described *diaschisis* as a distant effect of focal brain injury, characterized by an “abolition of excitability” and a resulting “functional standstill” in structurally intact but connected regions. This concept resonates closely with the disfacilitation mechanism explored here, where the loss of excitatory input from the lesioned area induces a leftward shift in the bifurcation diagram (Fig. 2), pushing the system toward a regime of SW activity^95^. In this framework, prolonged neuronal silence associated with SWs emerges around the lesion and propagates along anatomical pathways in a topology- and timing-dependent manner, shaping the spatial extent of remote functional alterations beyond the primary structural damage. These neuronal silences, which transiently interrupt communication within the cortical network—as observed during sleep^96^, anesthesia^97^, and in awake brain-injured patients^87^—have been associated with impaired behavioral performance in both rodents and humans^21–25,83^, thereby representing a plausible mechanism for the cognitive and behavioral impairments observed following focal brain lesions.

Importantly, this dynamic perspective on diaschisis complements static structural disconnectomics approaches^98^, which map lesion-induced white-matter disconnections to account for deficits extending beyond the visible injury. Functional magnetic resonance imaging (fMRI) studies further show that even small focal lesions can disrupt a substantial proportion of brain connections, resulting in widespread alterations in functional connectivity^99^. Building on these findings, our model suggests that post-lesional neuronal silencing, and its propagation throughout the connectome, may underlie these large-scale functional alterations, thereby bridging classical electrophysiological observations with modern fMRI findings. Supporting this view, recent evidence shows that enhanced slow/delta power is associated with the magnitude and spatial profile of fMRI connectivity alterations^100^, linking electrophysiological and hemodynamic markers of diaschisis and recovery.

Our framework can also be extended to other brain pathologies that exhibit pathological EEG slowing and behavioral deficits. Comparable slowing occurs in multiple sclerosis^101–106^, traumatic brain injury^107,108^, brain tumors^10^, and Parkinson’s disease^109^, where inflammation, white-matter lesions, or diffuse network disruption increase delta- and theta-band activity associated with cognitive impairment. Therefore, SW activity may represent a shared neurophysiological marker underlying functional deficits across diverse forms of brain damage.

In light of the results presented here, clear limitations and open avenues emerge. At the microscale, while our population model allowed us to investigate disconnection and spontaneous propagation patterns, it did not account for other mechanisms known to have a local impact on cortical dynamics following brain injury^11^. These include GABA-mediated excitation/inhibition imbalance^31,32,34^, cortical hyperexcitability^110,111^, local temperature increase^112^, inflammation^113,114^, and hypoxia ^115^. These factors could be addressed in future work and considered as additive contributors to perilesional SW generation, potentially enhancing SW propagation from the initial SW foci alongside the structural disconnection explored in our study. Further, our large-scale model can be tuned to subject-specific dynamics^116,46,47^ by incorporating personalized structural and functional connectivity derived from patient lesion data^117^. This approach would enable predictions about how specific lesions affect large-scale spontaneous^118^ and evoked^47^ dynamics.

Extending the factors of SW generation and propagation explored in our study, the model further suggests a potential role for arousal fluctuations^29,119–124^, which could be investigated in future work. Arousal, driven by brainstem and thalamic inputs, fluctuates over time and affects adaptation currents via potassium channel modulation^125,126^. Because disconnected regions lie closer to the bifurcation point for SW generation, the model predicts that fluctuations in adaptation (mimicking physiological arousal dynamics) that would have moderate effects in healthy brains could instead trigger lesion-specific topographical patterns of enhanced SW activity in the damaged brain.

Finally, our results also raise several questions regarding therapeutic strategies aimed at renormalizing the network-level alterations caused by pathological SWs. On one hand, locally confined SW activity around the injured site, identified in our model as first-degree disconnected nodes, may play adaptive or beneficial roles as it does during sleep^127,128^, and whether its dampening is desirable remains an open question. On the other hand, large-scale propagation of neuronal silence, which disrupts network communication, could potentially be mitigated by dampening or desynchronizing SW sources (i.e., by reducing SW coherence), as suggested by our modeling results (Fig. 5F). By simulating lesion-specific network dynamics, our model can be employed to predict how interventions—such as non-invasive brain stimulation techniques including repetitive TMS, transcranial alternating current stimulation, or temporal interference electrical stimulation^129–135^—might selectively reduce pathological SW propagation while preserving potentially beneficial local activity, thereby informing more targeted and effective therapeutic strategies.

## Acknowledgements

The authors thank Daniel Levenstein for support with the Wilson–Cowan model, and Marta Porro, Sasha D’Ambrosio, Ezequiel Mikulan, and Andrea Pigorini for their comments and suggestions on the manuscript. This work was financially supported by the following entities: ERC-2022-SYG Grant number 101071900 neurological mechanisms of injury and sleep-like cellular dynamics (NEMESIS); Italian National Recovery and Resilience Plan (NRRP), M4C2, funded by the European Union - NextGenerationEU (Project IR0000011, CUP B51E22000150006, “EBRAINS-Italy”); European Union’s Horizon 2020 Framework Program for Research and Innovation under the Specific Grant Agreement No.945539 (Human Brain Project SGA3); Tiny Blue Dot Foundation; Canadian Institute for Advanced Research (CIFAR), Canada; Italian Ministry for Universities and Research (PRIN 2022); Fondazione Regionale per la Ricerca Biomedica (Regione Lombardia), Project ERAPERMED2019–101, GA 779282; Project INFRASLOW PID2023-152918OB-I00 financed by MICIU / AEI / 10.13039/501100011033/FEDER, UE; Departament de Recerca i Universitats de la Generalitat de Catalunya (AGAUR 2021-SGR-01165 - NEUROVIRTUAL), supported by FEDER; Fondazione Cassa di Risparmio di Padova e Rovigo (CARIPARO) Grant Agreement number 55403; Ministry of Health, Italy (RF-2008 −12366899) Brain connectivity measured with high-density electroencephalography: a novel neurodiagnostic tool for stroke-NEUROCONN; BIAL foundation grant (Grant Agreement number 361/18); H2020 European School of Network Neuroscience (euSNN); H2020 Visionary Nature Based Actions For Heath, Wellbeing & Resilience in Cities (VARCITIES); Ministry of Health Italy (RF-2019-12369300): Eye-movement dynamics during free viewing as biomarker for assessment of visuospatial functions and for closed-loop rehabilitation in stroke (EYEMOVINSTROKE); Brazilian agency CNPQ (Grant Agreement number 444500/2024-3).

## 1. Supplementary Methods

### 1.1 Wilson-Cowan rate model with Spike-frequency adaptation

We implemented in The Virtual Brain (TVB) a Wilson-Cowan (WC) rate model with spike frequency adaptation current (Fig. S3A), as in Levenstein et al^1^. This model reproduces key features of Up and Down state transitions arising from the interplay between adaptation, recurrent excitation, external inputs, and noise. We used it to validate our main findings with a simplified rate-based model. The WC comprises three state variables representing the excitatory firing rate r_e_, inhibitory firing rate r_i_, and adaptation current *a*. Two independent Ornstein-Uhlenbeck (OU) noise processes, u_e_ and u_i_, act on the excitatory and inhibitory populations, respectively. The population firing rates evolve according to:

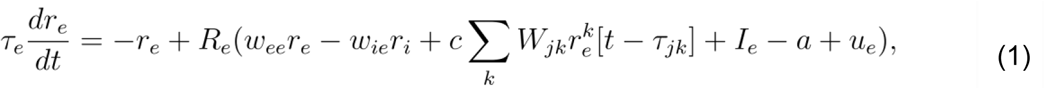

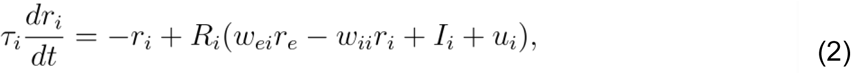

where R_e_ and R_i_ are power-law input-output functions^2^ defined as:

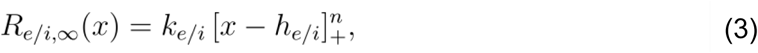

with scaling factors k_e_ and k_i_, thresholds h_e_ and h_i_, and exponent n. The internal coupling terms include recurrent weights w_ee_ (excitatory-to-excitatory), w_ei_ (excitatory-to-inhibitory), w_ie_ (inhibitory-to-excitatory), and w_ii_ (inhibitory-to-inhibitory neurons); the network coupling, which targets the excitatory population, is given by c∑_k_W_jk_r_e_^k^[t-𝜏_jk_], wher_e Wjk is the connec_tion weight from presynaptic node k to postsynaptic node j, and 𝜏_jk_ represents the time delay between j and k. The term c is the global coupling factor that scales all the connection weights. Thus, the effective excitatory coupling strength between j and k is given by the product of W_jk_ and c, and we denote it by C_jk_. r_e_^k^ is the firing rate of node k, and I_e_ and I_i_ are external inputs to excitatory and inhibitory populations, respectively. The slow adaptation current *a* implements spike-frequency adaptation through:

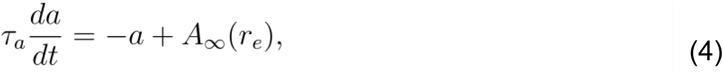

where

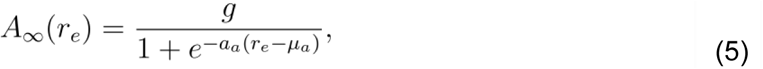

is a sigmoid function of the excitatory firing rate with gain a_a_, threshold μ_a_, and strength g (denoted as b in the original article). 𝜏_a_ is the time constant of the adaptation dynamics. The noise processes follow the OU dynamics:

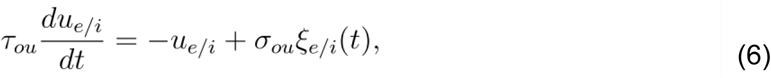

where ξ_e\i_(t) are independent Wiener processes with amplitude σ_ou_ and time constant 𝜏_ou_. The parameters used for the WC simulations are listed in Table S2. Equations were integrated using the Heun stochastic method in TVB with a time step of 0.5 ms.

### 1.2 Simulation and analysis of coupled WC models: from two nodes to whole-brain networks

#### Two-population model

To assess the effects of disconnectivity on the emergence of slow wave (SW) activity, a two-node model was implemented, consisting of two bidirectionally connected WC nodes (denoted WC_a_ and WC_b_) as in Section 2.3 of the main Methods. The nodes were coupled symmetrically or asymmetrically through excitatory-to-excitatory connections. The system behavior was explored parametrically as a function of the adaptation strength (g) and the excitatory coupling strength (C). In the symmetric configuration, g and C were varied equally in both populations, while in the asymmetric configuration, they were varied independently. For each parameter configuration, an independent 25s simulation was performed, with g varying from 0 to 4 in increments of 0.4, and C varying from 0 to 1 in increments of 0.05.

#### Whole-brain network model

The human whole-brain simulations were implemented using the same structural connectivity dataset described in the main Methods (Section 2.5). The connectivity matrix comprised 80 cortical regions from the Automatic Anatomical Labeling (AAL) atlas, averaged across subjects and thresholded to retain only connections present in more than 50% of individuals. The weights were normalized by in-degree to equalize excitatory input across nodes while preserving the structural topology. The model was first simulated in a control configuration, followed by lesion simulations in which each node was disconnected sequentially by setting its incoming and outgoing weights to zero. Each simulation lasted 100s, with a global coupling factor c=0.45 and g=1.8.

#### Spectral analysis

The power spectral density (psd) of the excitatory firing rate r_e_ was computed using Welch’s method with 3-s segments and 50% overlap. δ power (<4 Hz) was extracted from the psd and used as an index of SW activity. To characterize the SW activity impinging on a given node, two neighborhood-level measures were computed: i) Neighborhood δ power (NBR δ power): the average δ power across the afferent nodes projecting to a given node, ii) Neighborhood δ coherence (NBR δ coherence): the average pairwise spectral coherence in the δ band among afferent nodes of each target node. These measures are in accordance with the NBR %Down and NBR DownOverlap metrics described for the Jansen–Rit model (Section 2.6), but are computed in the spectral domain. The definition of the neighborhood was the same as that described in Section 2.6 of the main Methods.

#### Statistical analysis

Statistical analyses were performed as described in the main Methods (Section 2.9). The relationships between δ power (and its square root), connection loss, NBR δ power, and NBR δ coherence were assessed using Pearson’s correlation and Partial Least Squares (PLS) regression. As in the main analyses, the % variance explained (R^2^) was computed under both full and 10-fold cross-validation to evaluate model generalization. The contribution of each predictor to the first PLS component (PLS1) was quantified using Variable Importance in Projection (VIP) scores.

---

**Figure S1.**
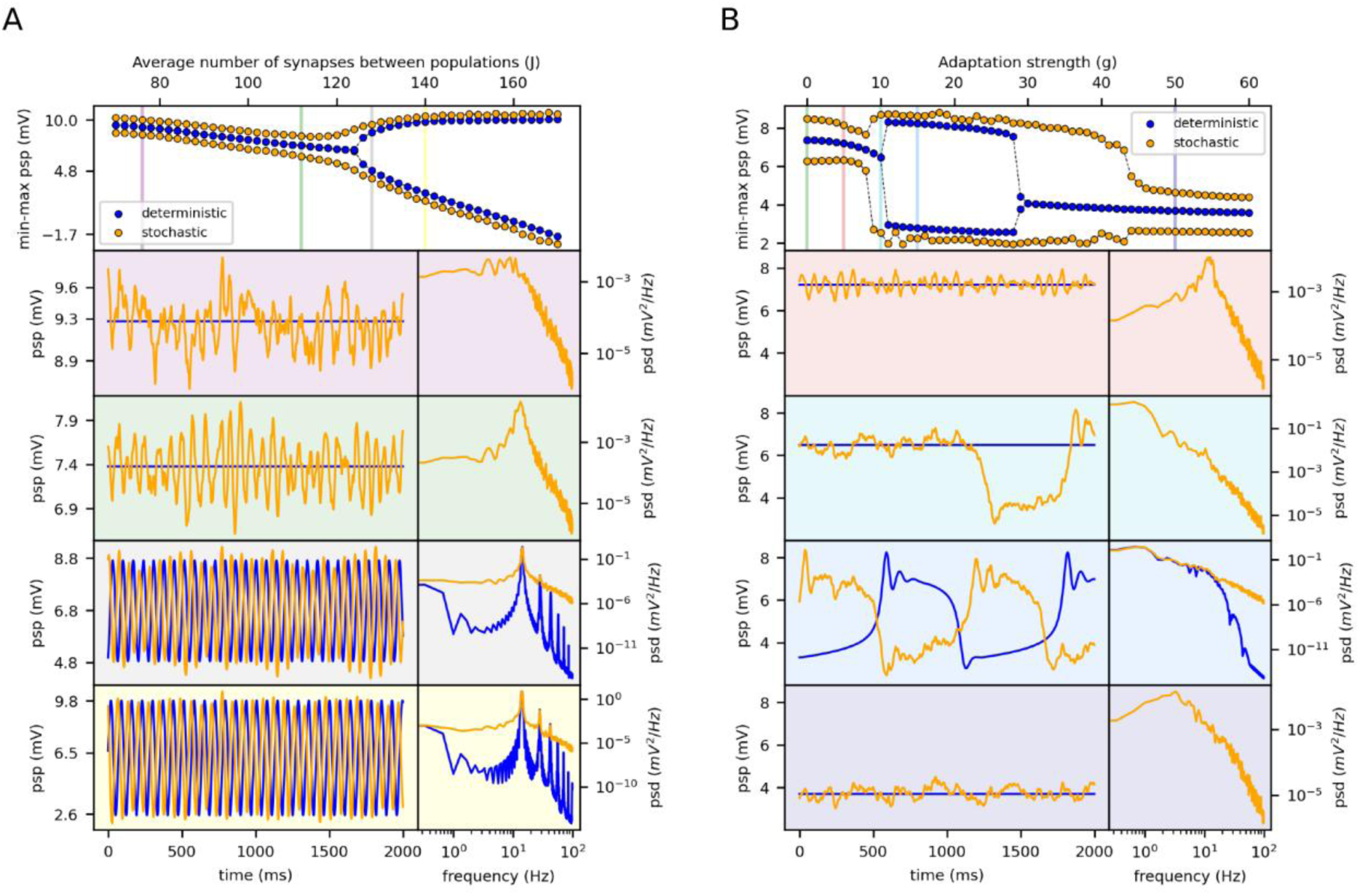
Bifurcation diagrams and oscillatory dynamics in deterministic and stochastic JR model. (A) Bifurcation diagram showing the min-max range of postsynaptic potential (PSP) as a function of J, for the deterministic (blue) and stochastic (yellow) models. Vertical bars mark specific values of J, for which representative psp traces (left) and their power spectral densities (psd, right) are shown. The background color of each panel corresponds to the respective vertical colored bar in the bifurcation diagram. The model was run without adaptation (g=0). A supercritical Hopf bifurcation occurs around J=128. (B) Bifurcation diagram as a function of adaptation strength (g), with the same conventions as in (A). The synaptic strength J is set to 112 (matching the green vertical bar in panel A). When g=0 the system corresponds to the same configuration of panel (A) for J=112. For low g, the system is near a supercritical Hopf bifurcation and exhibits damped oscillations. For g>9, Up/Down oscillations emerge. At g=9 and g=10, stochastic fluctuations drive transient transitions to the Down state. For g>39, the system stabilizes in a persistent Down state, with occasional Up state excursions for 30<g<39. Vertical bars indicate specific values of g (i.e., g=0, g=5, g=10, g=15, g=50), for which corresponding PSP traces and psd are shown in the lower panels, color-coded accordingly. Stability analysis of the deterministic model was performed in XPPAUT.

**Figure S2.**
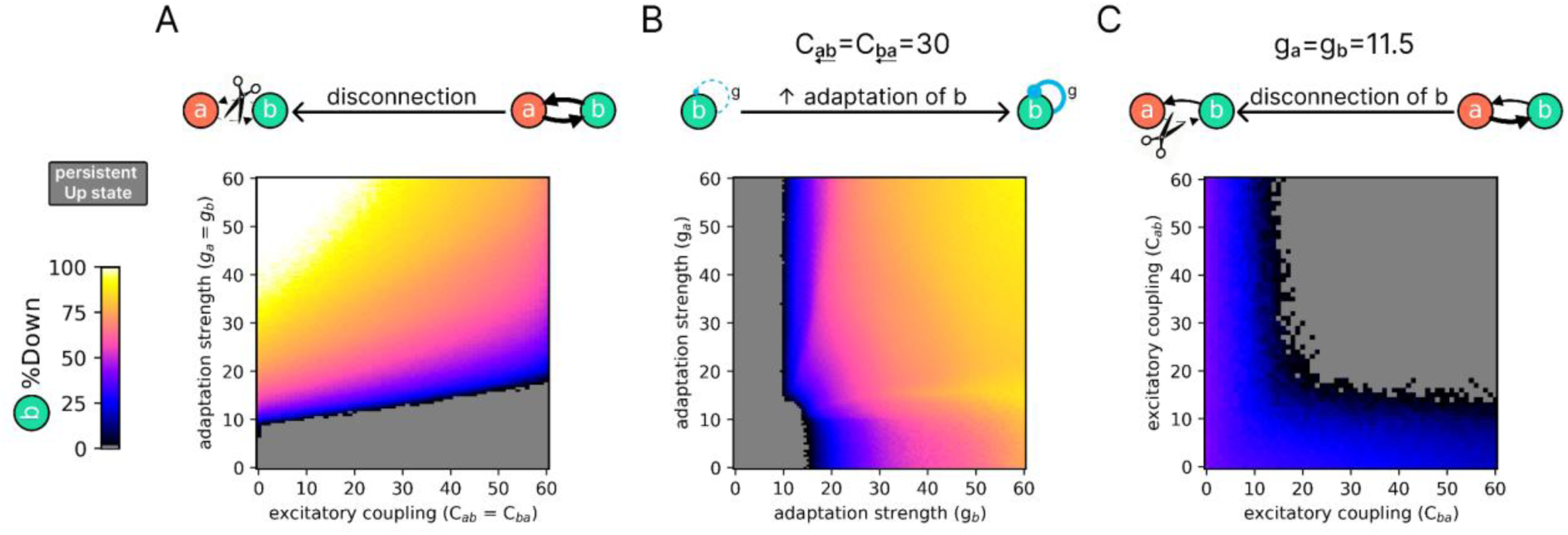
Effects of adaptation strength and excitatory coupling on slow wave emergence in population JR_b_. Each heatmap shows the percentage of time spent in the Down state (%Down) in JR_b_ as a function of (A) symmetric excitatory coupling (C_ab_ = C_ba_) and adaptation strength (g_a_ = g_b_), (B) asymmetric adaptation (g_a_, g_b_) with fixed symmetric coupling (C_ab_ = C_ba_ = 30), and (C) asymmetric coupling (C_ab_, C_ba_) with fixed adaptation strength (g_a_ = g_b_ = 11.5). The color scale indicates the proportion of time spent in the Down state, from persistent Up state (gray) to predominant Down state (white).

**Figure S3.**
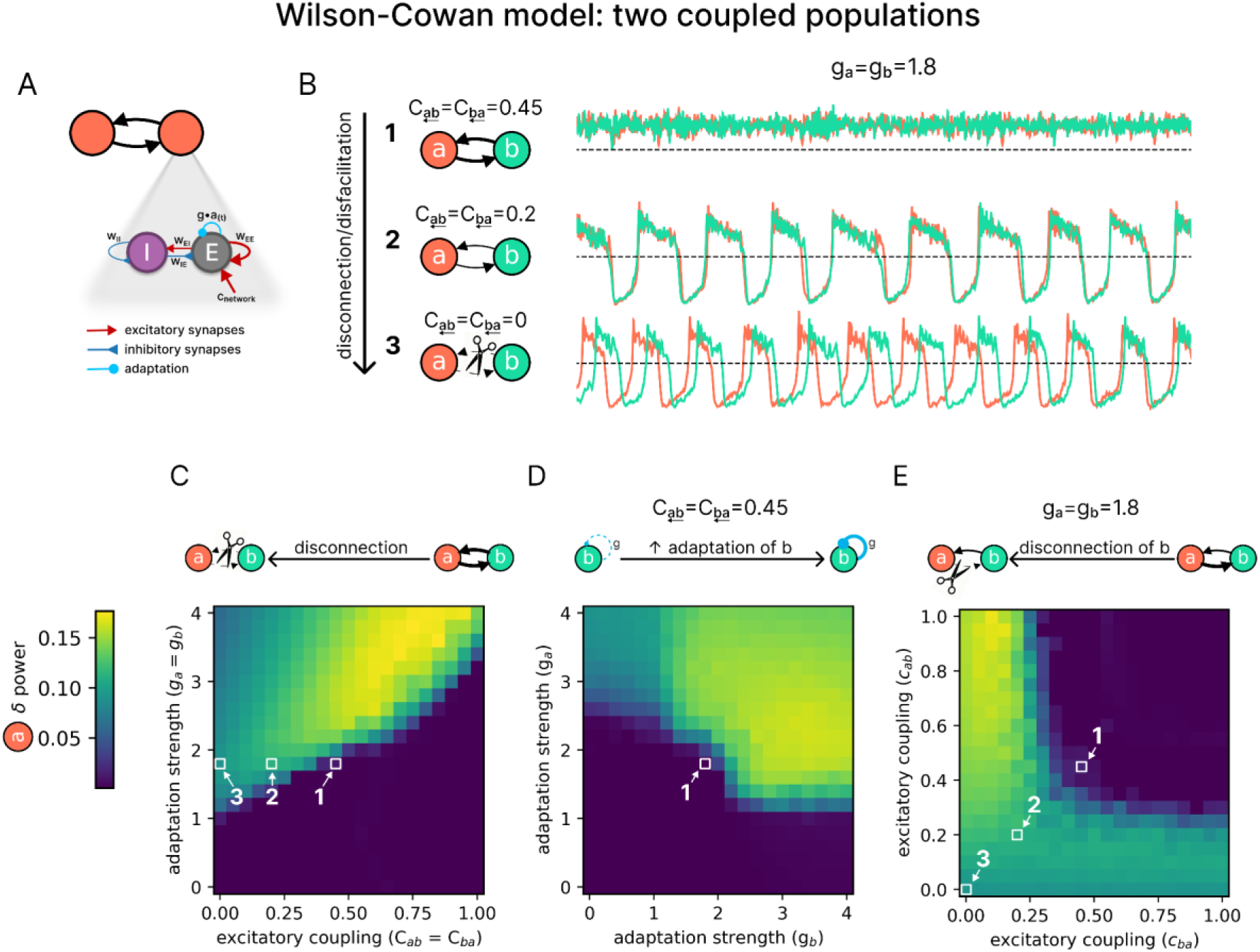
Effects of adaptation strength and excitatory coupling on slow wave emergence in two coupled WC nodes. (A) Representation of the Wilson-Cowan (WC) model with spike-frequency adaptation, consisting of interconnected excitatory (E) and inhibitory (I) neuronal pools. (B) Example dynamics of two symmetrically coupled populations (WC_a_ in orange, WC_b_ in cyan) for three levels of excitatory coupling (C_ab_=C_ba_). With C=0.45 and g=1.8 (top, #1), both populations show fast oscillations in a persistent Up state. Reducing coupling to C=0.2 (middle, #2) induces a transition to slow oscillations (SO). Complete disconnection (C=0, bottom, #3) further enhances SO activity.. (C) Heatmap showing the δ power of node WC_a_ as a function of adaptation strength (g, y-axis) and symmetric excitatory coupling (C, x-axis). The three squares with numbers correspond to the configuration of panel B, highlighting the effect of reducing C with g=0.45 kept fixed. Notice how the δ power increases through the transition from a persistent Up state (δ power) to SO (high δ power). (D) Effect of asymmetric adaptation. Heatmap shows δ power in WC_a_ as a function of g_b_ (x-axis) and g_a_ (y-axis), with symmetric coupling fixed at C=0.45. From the square #1, varying g_b_ (i.e., changing the adaptation of WC_b_) alone can induce a transition from a persistent Up state to SO (high δ power) in WC_a_, despite g_a_ being kept constant. (E) Effect of asymmetric coupling. Heatmap shows δ power in WC_a_ as a function of C_ba_ (x-axis) and C_ab_ (y-axis), with g=1.8 kept fixed. From the square #1, reducing C_ba_ (i.e., disconnecting WC_b_) alone is sufficient to induce a transition from a persistent Up state to SO (high δ power) in WC_a_, even when C_ab_ remains constant. Numbers 1, 2, and 3 in all panels correspond to identical parameter configurations across plots.

**Figure S4.**
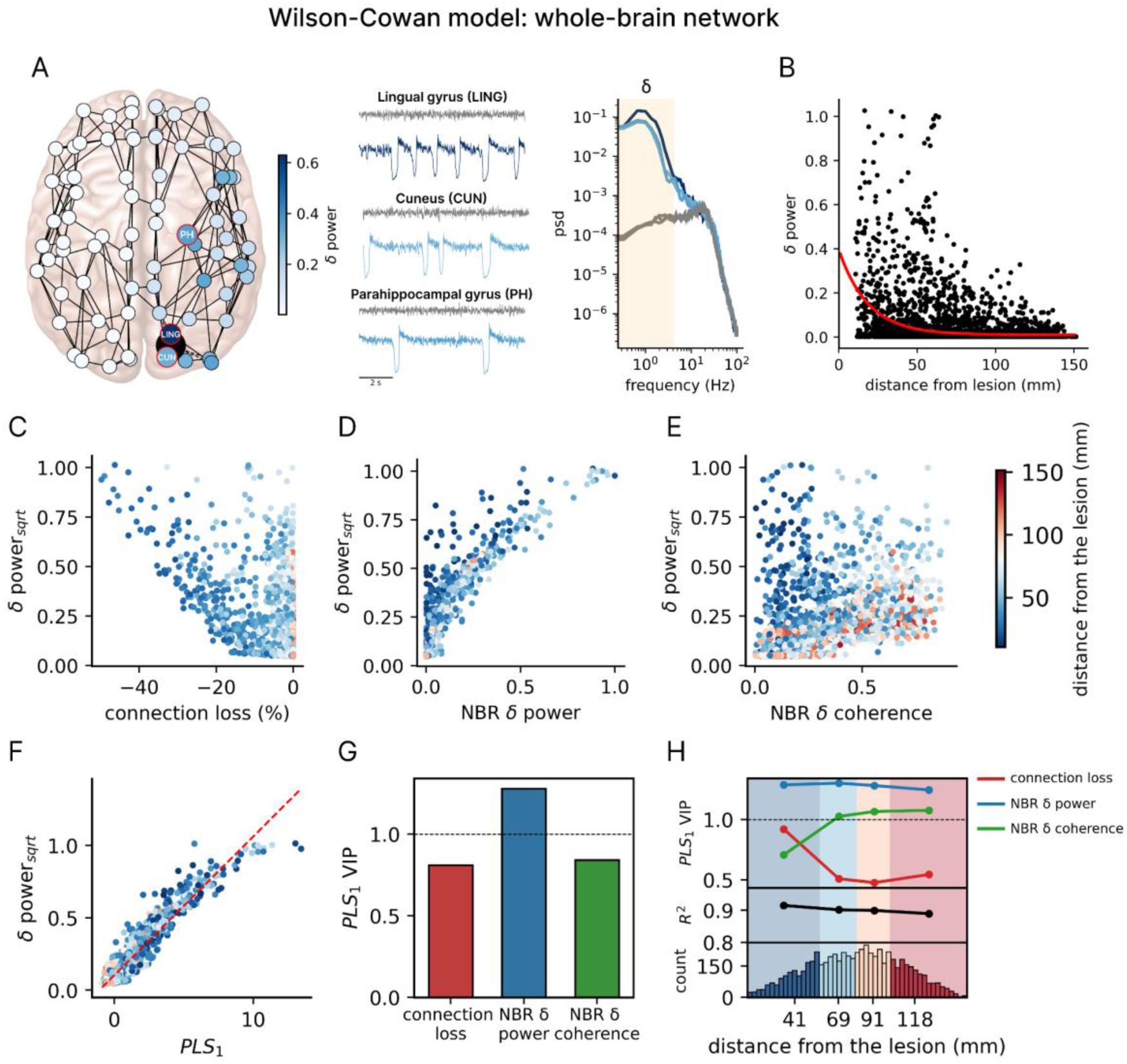
Slow wave generation and propagation following lesion in a whole-brain network of WC nodes. (A) Whole-brain network model with nodes defined by the Automatic Anatomical Labeling (AAL) atlas. The lesion of a node (here, the right calcarine cortex) is represented as a black dot, with the impact on the intact nodes quantified with δ power. The node colors indicate δ power, as shown in the accompanying color bar. Traces illustrate simulated activity from three representative regions in the pre-lesion (control; gray) and post-lesion (blue hues) conditions. In the control condition, the nodes display low-amplitude and fast oscillations in a persistent Up state (awake-like regime). After the lesion, SWs emerge along with an increase in δ power in the power spectral density (psd). The δ band is highlighted by the yellow span in the psd. (B) δ power as a function of the Euclidean distance from the lesion. The red curve represents an exponential fit (R^2^=0.11), indicating a distance-dependent decay in SW propagation. (C) Relationship between connection loss (percentage of lost coupling strength due to the lesion relative to control) and the square root of δ power (δ power_sqrt_). A negative relationship is observed (r=-0.57). (D) Relationship between the δ power of neighboring nodes (NBR δ power) and δ power_sqrt_. A positive relationship is observed (r=0.89). (E) Relationship between the coherence of SW activity of neighboring nodes (NBR δ coherence) and δ power_sqrt_. The variables show a positive relationship (r=0.59). (F) Partial least squares regression (PLS) of δ power_sqrt_ and the score of the first PLS component (PLS_1_) derived from the three independent variables (connection loss, NBR δ power, and NBR δ coherence). The model explained ∼90% of the variance (R^2^=0.92; after cross-validation R^2^=0.91). (G) PLS_1_ weights for the three independent variables, indicating their relative contributions. (H) Distance-dependent contributions of the three independent variables to δ power_sqrt_. Nodes were divided into quartiles based on their distance from the lesion, and PLS regression was performed for each subset. For each quartile, the model’s R^2^ and the VIP scores of each variable are reported.

**Table S1.**
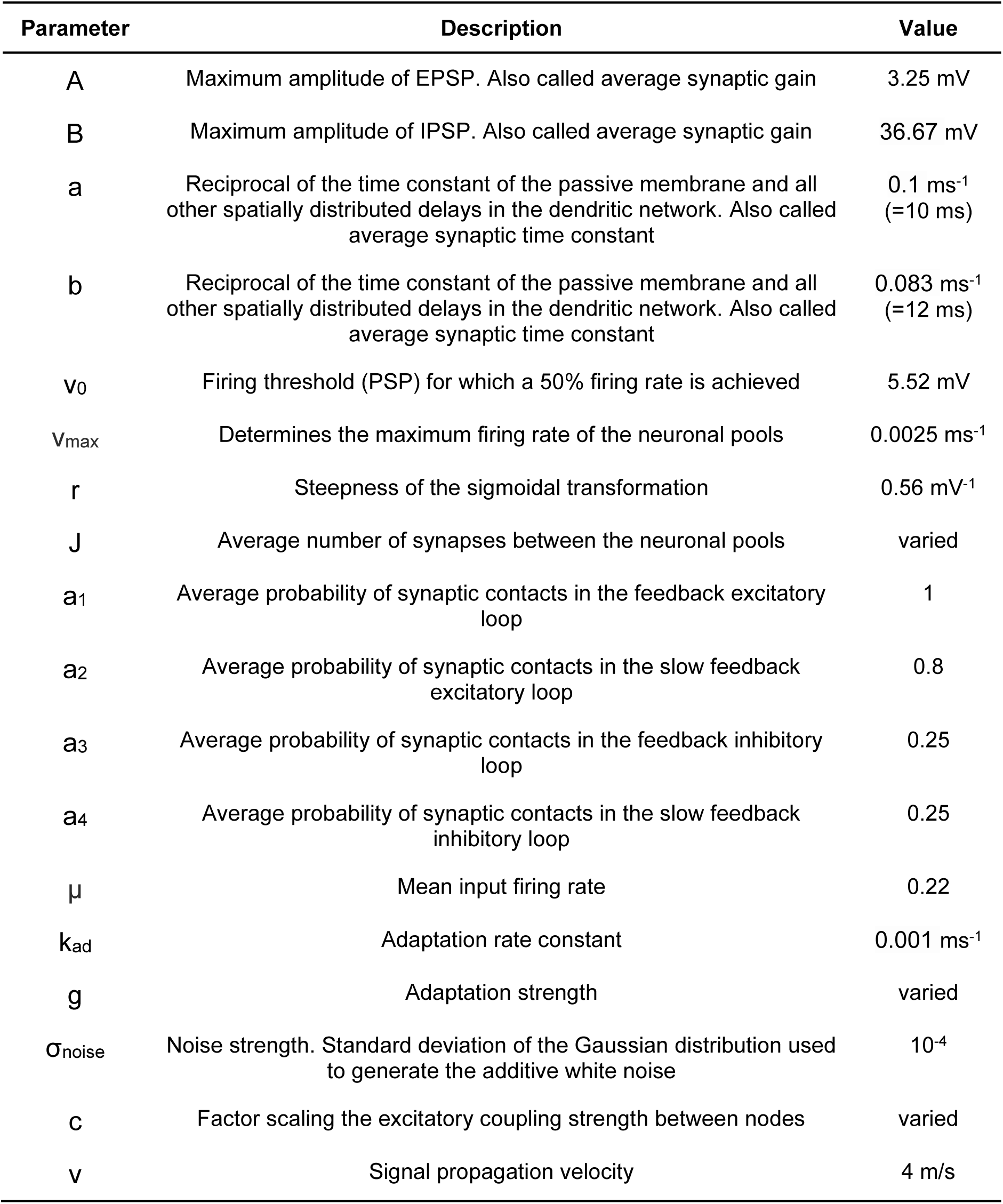
Parameters of the Jansen-Rit model used in the simulations. The table lists the parameters used in The Virtual Brain (TVB) implementation of the Jansen-Rit (JR) model with spike-frequency adaptation. All values lie within physiological ranges and have been previously explored in the literature (e.g., J and V ^3^; A, B, a, and b^4^; k_ad_^5^). The excitatory coupling strength factor c and the adaptation strength g were varied in the study and were set to c=30 and g=11.5 for both the toy and whole-brain networks. The parameter J was explored within the range used in^3^ and was set to produce subcritical dynamics just before a supercritical Hopf bifurcation, resulting in a peak in the power spectrum in the alpha band (J=121 for the whole-brain network model and J=112 for all other models).

**Table S2.**
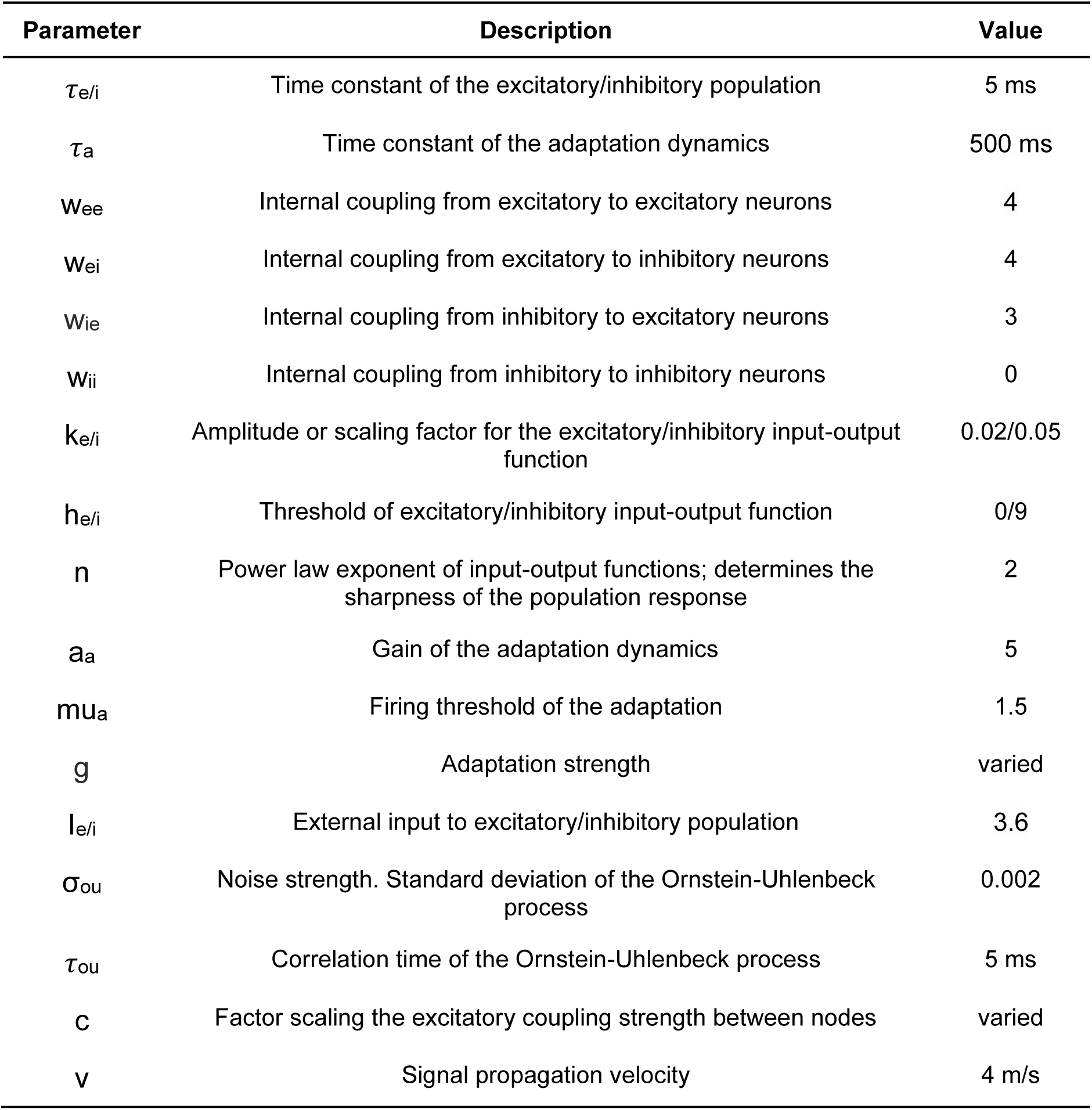
Parameters of the Wilson-Cowan model used in the simulations. The table lists the parameters used in The Virtual Brain (TVB) implementation of the rate model with spike-frequency adaptation. All parameter values were taken from the original publication^1^, except for the time constant of the adaptation dynamics (tau_a_, changed here from 200 to 500 ms to better align with the JR model while remaining consistent with adaptation kinetics^6^) and the threshold of the inhibitory input-output function (h_i_ changed from 12 to 9 to increase the contribution of the inhibitory population). In the two-coupled-population model, the adaptation strength (g; denoted as b in the original article) and the excitatory coupling strength (c) were varied, whereas in the whole-brain model, these were fixed to 1.8 and 0.45, respectively, to produce an awake-like regime in the intact model (pre-lesion).

## References

1. Carrera, E. & Tononi, G. Diaschisis: past, present, future. Brain 137, 2408–2422 (2014).

2. Fornito, A., Zalesky, A. & Breakspear, M. The connectomics of brain disorders. Nat Rev Neurosci 16, 159–172 (2015).

3. Baldassarre, A., Ramsey, L. E., Siegel, J. S., Shulman, G. L. & Corbetta, M. Brain connectivity and neurological disorders after stroke. Curr Opin Neurol 29, 706–713 (2016).

4. Grefkes, C., Eickhoff, S. B., Nowak, D. A., Dafotakis, M. & Fink, G. R. Dynamic intra-and interhemispheric interactions during unilateral and bilateral hand movements assessed with fMRI and DCM. NeuroImage 41, 1382–1394 (2008).

5. Siegel, J. S. et al. Disruptions of network connectivity predict impairment in multiple behavioral domains after stroke. Proceedings of the National Academy of Sciences 113, E4367–E4376 (2016).

6. Bottom-Tanzer, S. et al. Traumatic brain injury disrupts state-dependent functional cortical connectivity in a mouse model. Cereb Cortex 34, bhae038 (2024).

7. Latifi, S. & Carmichael, S. T. The emergence of multiscale connectomics-based approaches in stroke recovery. Trends in Neurosciences 47, 303–318 (2024).

8. von Monakow, C. Die Lokalisation im Grosshirn und der Abbau der Funktion durch Kortikale Herde. JAMA LXIII, 797 (1914).

9. Carrera, E. & Tononi, G. Diaschisis: past, present, future. Brain 137, 2408–2422 (2014).

10. Walter, W. G. The Electro-encephalogram in Cases of Cerebral Tumour. Proc R Soc Med 30, 579–598 (1937).

11. Massimini, M. et al. Sleep-like cortical dynamics during wakefulness and their network effects following brain injury. Nat Commun 15, 7207 (2024).

12. Steriade, M., Nunez, A. & Amzica, F. A novel slow (< 1 Hz) oscillation of neocortical neurons in vivo: depolarizing and hyperpolarizing components. J. Neurosci. 13, 3252–3265 (1993).

13. Sanchez-Vives, M. V. & McCormick, D. A. Cellular and network mechanisms of rhythmic recurrent activity in neocortex. Nat Neurosci 3, 1027–1034 (2000).

14. Luczak, A., Barthó, P., Marguet, S. L., Buzsáki, G. & Harris, K. D. Sequential structure of neocortical spontaneous activity in vivo. Proceedings of the National Academy of Sciences 104, 347–352 (2007).

15. Chauvette, S., Volgushev, M. & Timofeev, I. Origin of Active States in Local Neocortical Networks during Slow Sleep Oscillation. Cerebral Cortex 20, 2660–2674 (2010).

16. Nir, Y. et al. Regional Slow Waves and Spindles in Human Sleep. Neuron 70, 153–169 (2011).

17. Compte, A., Sanchez-Vives, M. V., McCormick, D. A. & Wang, X.-J. Cellular and Network Mechanisms of Slow Oscillatory Activity (<1 Hz) and Wave Propagations in a Cortical Network Model. Journal of Neurophysiology 89, 2707–2725 (2003).

18. Timofeev, I., Grenier, F. & Steriade, M. Disfacilitation and active inhibition in the neocortex during the natural sleep-wake cycle: An intracellular study. Proceedings of the National Academy of Sciences 98, 1924–1929 (2001).

19. Zucca, S. et al. An inhibitory gate for state transition in cortex. eLife 6, e26177 (2017).

20. Perez-Zabalza, M. et al. Modulation of cortical slow oscillatory rhythm by GABAB receptors: an in vitro experimental and computational study. J Physiol 598, 3439–3457 (2020).

21. Vyazovskiy, V. V. et al. Local sleep in awake rats. Nature 472, 443–447 (2011).

22. Bernardi, G. et al. Neural and Behavioral Correlates of Extended Training during Sleep Deprivation in Humans: Evidence for Local, Task-Specific Effects. J. Neurosci. 35, 4487–4500 (2015).

23. Nir, Y. et al. Selective neuronal lapses precede human cognitive lapses following sleep deprivation. Nat Med 23, 1474–1480 (2017).

24. Andrillon, T., Burns, A., Mackay, T., Windt, J. & Tsuchiya, N. Predicting lapses of attention with sleep-like slow waves. Nat Commun 12, 3657 (2021).

25. Marmelshtein, A., Eckerling, A., Hadad, B., Ben-Eliyahu, S. & Nir, Y. Sleep-like changes in neural processing emerge during sleep deprivation in early auditory cortex. Current Biology 33, 2925–2940.e6 (2023).

26. Gloor, P., Ball, G. & Schaul, N. Brain lesions that produce delta waves in the EEG. Neurology 27, 326–333 (1977).

27. Nuwer, M. R., Jordan, S. E. & Ahn, S. S. Evaluation of stroke using EEG frequency analysis and topographic mapping. Neurology 37, 1153–1159 (1987).

28. Meythaler, J. M., Peduzzi, J. D., Eleftheriou, E. & Novack, T. A. Current concepts: Diffuse axonal injury–associated traumatic brain injury. Archives of Physical Medicine and Rehabilitation 82, 1461–1471 (2001).

29. Schiff, N. D. Central Thalamic Contributions to Arousal Regulation and Neurological Disorders of Consciousness. Annals of the New York Academy of Sciences 1129, 105–118 (2008).

30. D’Ambrosio, S. et al. Detecting cortical reactivity alterations induced by structural disconnection in subcortical stroke. Clinical Neurophysiology 156, 1–3 (2023).

31. Imbrosci, B. & Mittmann, T. Functional Consequences of the Disturbances in the GABA-Mediated Inhibition Induced by Injuriesin the Cerebral Cortex. Neural Plast 2011, 614329 (2011).

32. Kim, Y. K., Yang, E. J., Cho, K., Lim, J. Y. & Paik, N.-J. Functional Recovery After Ischemic Stroke Is Associated With Reduced GABAergic Inhibition in the Cerebral Cortex: A GABA PET Study. Neurorehabil Neural Repair 28, 576–583 (2014).

33. Funk, C. M. et al. Role of Somatostatin-Positive Cortical Interneurons in the Generation of Sleep Slow Waves. J. Neurosci. 37, 9132–9148 (2017).

34. Santos, F. P. dos, Vohryzek, J. & Verschure, P. F. M. J. Multiscale effects of excitatory-inhibitory homeostasis in lesioned cortical networks: A computational study. PLOS Computational Biology 19, e1011279 (2023).

35. Russo, S. et al. Focal lesions induce large-scale percolation of sleep-like intracerebral activity in awake humans. NeuroImage 234, 117964 (2021).

36. Deco, G., Jirsa, V. K., Robinson, P. A., Breakspear, M. & Friston, K. The Dynamic Brain: From Spiking Neurons to Neural Masses and Cortical Fields. PLOS Computational Biology 4, e1000092 (2008).

37. Breakspear, M. Dynamic models of large-scale brain activity. Nat Neurosci 20, 340–352 (2017).

38. Traub, R. D. & Wong, R. K. Cellular mechanism of neuronal synchronization in epilepsy. Science 216, 745–747 (1982).

39. Jirsa, V. K. et al. The Virtual Epileptic Patient: Individualized whole-brain models of epilepsy spread. NeuroImage 145, 377–388 (2017).

40. Proix, T., Bartolomei, F., Guye, M. & Jirsa, V. K. Individual brain structure and modelling predict seizure propagation. Brain 140, 641–654 (2017).

41. Rigney, G., Lennon, M. & Holderrieth, P. The use of computational models in the management and prognosis of refractory epilepsy: A critical evaluation. Seizure 91, 132–140 (2021).

42. Jansen, B. H. & Rit, V. G. Electroencephalogram and visual evoked potential generation in a mathematical model of coupled cortical columns. Biol Cybern 73, 357–366 (1995).

43. David, O. & Friston, K. J. A neural mass model for MEG/EEG:: coupling and neuronal dynamics. NeuroImage 20, 1743–1755 (2003).

44. Coronel-Oliveros, C., Gießing, C., Medel, V., Cofré, R. & Orio, P. Whole-brain modeling explains the context-dependent effects of cholinergic neuromodulation. NeuroImage 265, 119782 (2023).

45. Sanchez-Todo, R. et al. A physical neural mass model framework for the analysis of oscillatory generators from laminar electrophysiological recordings. NeuroImage 270, 119938 (2023).

46. Momi, D., Wang, Z. & Griffiths, J. D. TMS-evoked responses are driven by recurrent large-scale network dynamics. Elife 12, e83232 (2023).

47. Momi, D. et al. Stimulation mapping and whole-brain modeling reveal gradients of excitability and recurrence in cortical networks. Nat Commun 16, 3222 (2025).

48. Koller, D. P., Schirner, M. & Ritter, P. Human connectome topology directs cortical traveling waves and shapes frequency gradients. Nat Commun 15, 3570 (2024).

49. Camassa, A., Galluzzi, A., Mattia, M. & Sanchez-Vives, M. V. Deterministic and Stochastic Components of Cortical Down States: Dynamics and Modulation. J. Neurosci. 42, 9387–9400 (2022).

50. Moran, R. J. et al. A neural mass model of spectral responses in electrophysiology. Neuroimage 37, 706–720 (2007).

51. Benda, J. & Herz, A. V. M. A universal model for spike-frequency adaptation. Neural Comput 15, 2523–2564 (2003).

52. Ermentrout, B. XPPAUT. in Computational Systems Neurobiology (ed. Le Novère, N.) 519–531 (Springer Netherlands, Dordrecht, 2012). doi:10.1007/978-94-007-3858-4_17.

53. Lemaréchal, J.-D. et al. A brain atlas of axonal and synaptic delays based on modelling of cortico-cortical evoked potentials. Brain 145, 1653–1667 (2021).

54. Škoch, A. et al. Human brain structural connectivity matrices–ready for modelling. Sci Data 9, 486 (2022).

55. Tzourio-Mazoyer, N. et al. Automated Anatomical Labeling of Activations in SPM Using a Macroscopic Anatomical Parcellation of the MNI MRI Single-Subject Brain. NeuroImage 15, 273–289 (2002).

56. de Reus, M. A. & van den Heuvel, M. P. Estimating false positives and negatives in brain networks. NeuroImage 70, 402–409 (2013).

57. Virtanen, P. et al. SciPy 1.0: fundamental algorithms for scientific computing in Python. Nat Methods 17, 261–272 (2020).

58. Wold, S., Johansson, E. & Cocchi, M. PLS: partial least squares projections to latent structures. in 3D QSAR in Drug Design: Theory, Methods and Applications. 523–550 (Kluwer ESCOM Science Publisher, 1993).

59. Wold, S., Sjöström, M. & Eriksson, L. PLS-regression: a basic tool of chemometrics. Chemometrics and Intelligent Laboratory Systems 58, 109–130 (2001).

60. Pedregosa, F. et al. Scikit-learn: Machine Learning in Python. J. Mach. Learn. Res. 12, 2825–2830 (2011).

61. Mehmood, T., Liland, K. H., Snipen, L. & Sæbø, S. A review of variable selection methods in Partial Least Squares Regression. Chemometrics and Intelligent Laboratory Systems 118, 62–69 (2012).

62. Gast, R. et al. PyRates—A Python framework for rate-based neural simulations. PLOS ONE 14, e0225900 (2019).

63. Linkenkaer-Hansen, K., Nikouline, V. V., Palva, J. M. & Ilmoniemi, R. J. Long-Range Temporal Correlations and Scaling Behavior in Human Brain Oscillations. J. Neurosci. 21, 1370–1377 (2001).

64. Destexhe, A., Hughes, S. W., Rudolph, M. & Crunelli, V. Are corticothalamic ‘up’ states fragments of wakefulness? Trends Neurosci 30, 334–342 (2007).

65. Compte, A. et al. Spontaneous High-Frequency (10–80 Hz) Oscillations during Up States in the Cerebral Cortex In Vitro. J. Neurosci. 28, 13828–13844 (2008).

66. Massimini, M., Huber, R., Ferrarelli, F., Hill, S. & Tononi, G. The Sleep Slow Oscillation as a Traveling Wave. J. Neurosci. 24, 6862–6870 (2004).

67. Torao-Angosto, M., Manasanch, A., Mattia, M. & Sanchez-Vives, M. V. Up and Down States During Slow Oscillations in Slow-Wave Sleep and Different Levels of Anesthesia. Front. Syst. Neurosci. 15, (2021).

68. Mattia, M. & Sanchez-Vives, M. V. Exploring the spectrum of dynamical regimes and timescales in spontaneous cortical activity. Cogn Neurodyn 6, 239–250 (2012).

69. Cattani, A. et al. Adaptation Shapes Local Cortical Reactivity: From Bifurcation Diagram and Simulations to Human Physiological and Pathological Responses. eNeuro 10, (2023).

70. Timofeev, I., Grenier, F., Bazhenov, M., Sejnowski, T. J. & Steriade, M. Origin of slow cortical oscillations in deafferented cortical slabs. Cereb Cortex 10, 1185–1199 (2000).

71. Sarasso, S. et al. Local sleep-like cortical reactivity in the awake brain after focal injury. Brain 143, 3672–3684 (2020).

72. Tscherpel, C. et al. Local neuronal sleep after stroke: The role of cortical bistability in brain reorganization. Brain Stimulation: Basic, Translational, and Clinical Research in Neuromodulation 17, 836–846 (2024).

73. McCormick, D. A., Nestvogel, D. & He, B. J. Neuromodulation of Brain State and Behavior. Annu Rev Neurosci 43, 391–415 (2020).

74. Schwindt, P. C., Spain, W. J. & Crill, W. E. Long-lasting reduction of excitability by a sodium-dependent potassium current in cat neocortical neurons. J Neurophysiol 61, 233–244 (1989).

75. McCormick, D. A. & Williamson, A. Convergence and divergence of neurotransmitter action in human cerebral cortex. Proceedings of the National Academy of Sciences 86, 8098–8102 (1989).

76. Huber, R., Felice Ghilardi, M., Massimini, M. & Tononi, G. Local sleep and learning. Nature 430, 78–81 (2004).

77. Stickgold, R. Sleep-dependent memory consolidation. Nature 437, 1272–1278 (2005).

78. Tononi, G. & Cirelli, C. Sleep function and synaptic homeostasis. Sleep Med Rev 10, 49–62 (2006).

79. Marshall, L., Helgadóttir, H., Mölle, M. & Born, J. Boosting slow oscillations during sleep potentiates memory. Nature 444, 610–613 (2006).

80. Uji, M. et al. Human deep sleep facilitates cerebrospinal fluid dynamics linked to spontaneous brain oscillations and neural events. Proceedings of the National Academy of Sciences 122, e2509626122 (2025).

81. Butz, M. et al. Perilesional pathological oscillatory activity in the magnetoencephalogram of patients with cortical brain lesions. Neuroscience Letters 355, 93–96 (2004).

82. Sarasso, S. et al. Reduction of sleep-like perilesional slow waves and clinical evolution after stroke: A TMS-EEG study. Clinical Neurophysiology 175, 2110746 (2025).

83. Sheybani, L. et al. Wake slow waves in focal human epilepsy impact network activity and cognition. Nat Commun 14, 7397 (2023).

84. Lanzone, J. et al. EEG spectral exponent as a synthetic index for the longitudinal assessment of stroke recovery. Clin Neurophysiol 137, 92–101 (2022).

85. Covelo, J. et al. Spatiotemporal network dynamics and structural correlates in the human cerebral cortex *in vitro*. Progress in Neurobiology 246, 102719 (2025).

86. Colombo, M. A. et al. Hemispherotomy leads to persistent sleep-like slow waves in the isolated cortex of awake humans. PLOS Biology 23, e3003060 (2025).

87. Rosanova, M. et al. Sleep-like cortical OFF-periods disrupt causality and complexity in the brain of unresponsive wakefulness syndrome patients. Nat Commun 9, 4427 (2018).

88. Colombo, M. A. et al. Beyond alpha power: EEG spatial and spectral gradients robustly stratify disorders of consciousness. Cereb Cortex 33, 7193–7210 (2023).

89. Porta, L. D., Barbero-Castillo, A., Sanchez-Sanchez, J. M. & Sanchez-Vives, M. V. M-current modulation of cortical slow oscillations: Network dynamics and computational modeling. PLOS Computational Biology 19, e1011246 (2023).

90. Douglas, R. J. & Martin, K. A. C. Neuronal circuits of the neocortex. Annu Rev Neurosci 27, 419–451 (2004).

91. Bullmore, E. & Sporns, O. Complex brain networks: graph theoretical analysis of structural and functional systems. Nat Rev Neurosci 10, 186–198 (2009).

92. Bassett, D. S. & Sporns, O. Network neuroscience. Nat Neurosci 20, 353–364 (2017).

93. Boucsein, C., Nawrot, M., Schnepel, P. & Aertsen, A. Beyond the Cortical Column: Abundance and Physiology of Horizontal Connections Imply a Strong Role for Inputs from the Surround. Front. Neurosci. 5, (2011).

94. Ercsey-Ravasz, M. et al. A Predictive Network Model of Cerebral Cortical Connectivity Based on a Distance Rule. Neuron 80, 184–197 (2013).

95. Rabuffo, G. et al. Mapping global brain reconfigurations following local targeted manipulations. Proceedings of the National Academy of Sciences 122, e2405706122 (2025).

96. Pigorini, A. et al. Bistability breaks-off deterministic responses to intracortical stimulation during non-REM sleep. Neuroimage 112, 105–113 (2015).

97. Krom, A. J. et al. Anesthesia-induced loss of consciousness disrupts auditory responses beyond primary cortex. Proceedings of the National Academy of Sciences 117, 11770–11780 (2020).

98. Thiebaut de Schotten, M., Foulon, C. & Nachev, P. Brain disconnections link structural connectivity with function and behaviour. Nat Commun 11, 5094 (2020).

99. Griffis, J. C., Metcalf, N. V., Corbetta, M. & Shulman, G. L. Structural Disconnections Explain Brain Network Dysfunction after Stroke. Cell Reports 28, 2527–2540.e9 (2019).

100. Rocchi, F. et al. Increased fMRI connectivity upon chemogenetic inhibition of the mouse prefrontal cortex. Nat Commun 13, 1056 (2022).

101. Kassubek, J., Sörös, P., Kober, H., Stippich, C. & Vieth, J. B. Focal slow and beta brain activity in patients with multiple sclerosis revealed by magnetoencephalography. Brain Topogr 11, 193–200 (1999).

102. Van der Meer, M. L. et al. Cognition in MS correlates with resting-state oscillatory brain activity: An explorative MEG source-space study. Neuroimage Clin 2, 727–734 (2013).

103. Keune, P. M. et al. Frontal brain activity and cognitive processing speed in multiple sclerosis: An exploration of EEG neurofeedback training. NeuroImage: Clinical 22, 101716 (2019).

104. Krupina, N. A., Churyukanov, M. V., Kukushkin, M. L. & Yakhno, N. N. Central Neuropathic Pain and Profiles of Quantitative Electroencephalography in Multiple Sclerosis Patients. Front Neurol 10, 1380 (2020).

105. Salim, A. A., Ali, S. H., Hussain, A. M. & Ibrahim, W. N. Electroencephalographic evidence of gray matter lesions among multiple sclerosis patients. Medicine (Baltimore) 100, e27001 (2021).

106. Mórocz, I., et al. Brain states analysis of EEG predicts multiple sclerosis and mirrors disease duration and burden. Preprint at 10.48550/arXiv.2406.15665 (2025).

107. Mofakham, S. et al. Electrocorticography reveals thalamic control of cortical dynamics following traumatic brain injury. Commun Biol 4, 1210 (2021).

108. Tewarie, P. K. B. et al. Early EEG monitoring predicts clinical outcome in patients with moderate to severe traumatic brain injury. Neuroimage Clin 37, 103350 (2023).

109. Bernasconi, F. et al. Sleep-like slow waves during wakefulness uncover a malignant form of Parkinson’s disease. 2025.08.29.25334276 Preprint at 10.1101/2025.08.29.25334276 (2025).

110. Schiene, K. et al. Neuronal Hyperexcitability and Reduction of GABAA-Receptor Expression in the Surround of Cerebral Photothrombosis. J Cereb Blood Flow Metab 16, 906–914 (1996).

111. Koch, G. et al. Hyperexcitability of parietal-motor functional connections for the intact left-hemisphere in neglect patients. Brain 131, 3147–3155 (2008).

112. Karaszewski, B. et al. Measurement of brain temperature with magnetic resonance spectroscopy in acute ischemic stroke. Annals of Neurology 60, 438–446 (2006).

113. Yoshida, H. et al. State-specific asymmetries in EEG slow wave activity induced by local application of TNFα. Brain Research 1009, 129–136 (2004).

114. Yasuda, T., Yoshida, H., Garcia-Garcia, F., Kay, D. & Krueger, J. M. Interleukin-1β has a Role in Cerebral Cortical State-Dependent Electroencephalographic Slow-Wave Activity. Sleep 28, 177–186 (2005).

115. Sun, H. & Feng, Z. Neuroprotective role of ATP-sensitive potassium channels in cerebral ischemia. Acta Pharmacol Sin 34, 24–32 (2013).

116. Capone, C. et al. Simulations approaching data: cortical slow waves in inferred models of the whole hemisphere of mouse. Commun Biol 6, 1–14 (2023).

117. Idesis, S. et al. Generative whole-brain dynamics models from healthy subjects predict functional alterations in stroke at the level of individual patients. Brain Communications 6, fcae237 (2024).

118. Idesis, S. et al. Whole-brain model replicates sleep-like slow-wave dynamics generated by stroke lesions. Neurobiology of Disease 200, 106613 (2024).

119. Makeig, S. & Inlow, M. Lapse in alertness: coherence of fluctuations in performance and EEG spectrum. Electroencephalography and Clinical Neurophysiology 86, 23–35 (1993).

120. Makeig, S. & Jung, T. P. Changes in alertness are a principal component of variance in the EEG spectrum. Neuroreport 7, 213–216 (1995).

121. Berridge, C. W. Noradrenergic modulation of arousal. Brain Research Reviews 58, 1–17 (2008).

122. Raut, R. V. et al. Global waves synchronize the brain’s functional systems with fluctuating arousal. Science Advances 7, eabf2709 (2021).

123. Raut, R. V. et al. Arousal as a universal embedding for spatiotemporal brain dynamics. Nature 1–8 (2025) doi:10.1038/s41586-025-09544-4.

124. Carro-Domínguez, M. et al. Pupil size reveals arousal level fluctuations in human sleep. Nat Commun 16, 2070 (2025).

125. McCormick, D. A. Neurotransmitter actions in the thalamus and cerebral cortex and their role in neuromodulation of thalamocortical activity. Prog Neurobiol 39, 337–388 (1992).

126. Lee, S.-H. & Dan, Y. Neuromodulation of Brain States. Neuron 76, 209–222 (2012).

127. Carmichael, S. T. & Chesselet, M.-F. Synchronous Neuronal Activity Is a Signal for Axonal Sprouting after Cortical Lesions in the Adult. J. Neurosci. 22, 6062–6070 (2002).

128. Krone, L. B. & Vyazovskiy, V. V. Unresponsive or just asleep? Do local slow waves in the perilesional cortex have a function? Brain 143, 3513–3515 (2020).

129. Xie, Y. & Zhang, T. Repetitive transcranial magnetic stimulation improves consciousness disturbance in stroke patients: A quantitative electroencephalography spectral power analysis. Neural Regen Res 7, 2465–2472 (2012).

130. Garside, P., Arizpe, J., Lau, C.-I., Goh, C. & Walsh, V. Cross-hemispheric Alternating Current Stimulation During a Nap Disrupts Slow Wave Activity and Associated Memory Consolidation. Brain Stimul 8, 520–527 (2015).

131. Grossman, N. et al. Noninvasive Deep Brain Stimulation via Temporally Interfering Electric Fields. Cell 169, 1029–1041.e16 (2017).

132. Fehér, K. D. et al. Shaping the slow waves of sleep: A systematic and integrative review of sleep slow wave modulation in humans using non-invasive brain stimulation. Sleep Medicine Reviews 58, 101438 (2021).

133. Violante, I. R. et al. Non-invasive temporal interference electrical stimulation of the human hippocampus. Nat Neurosci 26, 1994–2004 (2023).

134. Vieira, P. G., Krause, M. R. & Pack, C. C. Temporal interference stimulation disrupts spike timing in the primate brain. Nat Commun 15, 4558 (2024).

135. Covelo, J., Cortada, M., Vinci, G. V., Mattia, M. & Sanchez-Vives, M. V. Network Desynchronization with Sine Waves: from Synchrony to Asynchrony by Periodic Stimulation. Advanced Science 12, e14602 (2025).

## Supplementary References

1. Levenstein, D., Buzsáki, G. & Rinzel, J. NREM sleep in the rodent neocortex and hippocampus reflects excitable dynamics. Nat Commun 10, 2478 (2019).

2. Ahmadian, Y., Rubin, D. B. & Miller, K. D. Analysis of the stabilized supralinear network. Neural Comput 25, 1994–2037 (2013).

3. Jansen, B. H. & Rit, V. G. Electroencephalogram and visual evoked potential generation in a mathematical model of coupled cortical columns. Biol Cybern 73, 357–366 (1995).

4. David, O. & Friston, K. J. A neural mass model for MEG/EEG: coupling and neuronal dynamics. NeuroImage 20, 1743–1755 (2003).

5. Moran, R. J. et al. A neural mass model of spectral responses in electrophysiology. Neuroimage 37, 706–720 (2007).

6. Benda, J. & Herz, A. V. M. A universal model for spike-frequency adaptation. Neural Comput 15, 2523–2564 (2003).

