## Supplementary material for "Slow waves generation and propagation in a model of brain lesions"

### Supplementary Information

#### Table of Contents

**1. Supplementary Methods**
- Wilson-Cowan rate model with spike frequency adaptation
- Simulation and analysis of coupled WC rate models: from two nodes to whole-brain networks

**2. Supplementary Figures**
- Figure S1. Bifurcation diagrams and oscillatory dynamics in the deterministic and stochastic JR model
- Figure S2. Effects of adaptation strength and excitatory coupling on slow wave emergence in population $JR_b$
- Figure S3. Effects of adaptation strength and excitatory coupling on slow wave emergence in two coupled WC nodes
- Figure S4. Slow wave generation and propagation following lesion in a whole-brain network of WC nodes

**3. Supplementary Tables**
- Table S1. Parameters of the Jansen-Rit model used in the simulations
- Table S2. Parameters of the WC model used in the simulations

**4. Supplementary References**

### 1. Supplementary Methods

#### 1.1 Wilson-Cowan rate model with Spike-frequency adaptation

We implemented in The Virtual Brain (TVB) a Wilson-Cowan (WC) rate model with spike frequency adaptation current (Fig. S3A), as in Levenstein et al[1]. This model reproduces key features of Up and Down state transitions arising from the interplay between adaptation, recurrent excitation, external inputs, and noise. We used it to validate our main findings with a simplified rate-based model. The WC comprises three state variables representing the excitatory firing rate $r_e$, inhibitory firing rate $r_i$, and adaptation current *a*. Two independent Ornstein-Uhlenbeck (OU) noise processes, $u_e$ and $u_i$, act on the excitatory and inhibitory populations, respectively. The population firing rates evolve according to:

$$\tau_e \frac{dr_e}{dt} = -r_e + R_e(w_{ee}r_e - w_{ie}r_i + c\sum_k W_{jk} r_e^k[t-\tau_{jk}] + I_e - a + u_e), \quad (1)$$

$$\tau_i \frac{dr_i}{dt} = -r_i + R_i(w_{ei}r_e - w_{ii}r_i + I_i + u_i), \quad (2)$$

where $R_e$ and $R_i$ are power-law input-output functions[2] defined as:

$$R_{e/i,\infty}(x) = k_{e/i}\,[x - h_{e/i}]_+^n, \quad (3)$$

with scaling factors $k_e$ and $k_i$, thresholds $h_e$ and $h_i$, and exponent n. The internal coupling terms include recurrent weights $w_{ee}$ (excitatory-to-excitatory), $w_{ei}$ (excitatory-to-inhibitory), $w_{ie}$ (inhibitory-to-excitatory), and $w_{ii}$ (inhibitory-to-inhibitory neurons); the network coupling, which targets the excitatory population, is given by $c\sum_k W_{jk} r_e^k[t-\tau_{jk}]$, where $W_{jk}$ is the connection weight from presynaptic node k to postsynaptic node j, and $\tau_{jk}$ represents the time delay between j and k. The term c is the global coupling factor that scales all the connection weights. Thus, the effective excitatory coupling strength between j and k is given by the product of $W_{jk}$ and c, and we denote it by $C_{jk}$. $r_e^k$ is the firing rate of node k, and $I_e$ and $I_i$ are external inputs to excitatory and inhibitory populations, respectively. The slow adaptation current *a* implements spike-frequency adaptation through:

$$\tau_a \frac{da}{dt} = -a + A_\infty(r_e), \quad (4)$$

where

$$A_\infty(r_e) = \frac{g}{1 + e^{-a_a(r_e - \mu_a)}}, \quad (5)$$

is a sigmoid function of the excitatory firing rate with gain $a_a$, threshold $\mu_a$, and strength g (denoted as b in the original article). $\tau_a$ is the time constant of the adaptation dynamics. The noise processes follow the OU dynamics:

$$\tau_{ou} \frac{du_{e/i}}{dt} = -u_{e/i} + \sigma_{ou}\xi_{e/i}(t), \quad (6)$$

where $\xi_{e\backslash i}(t)$ are independent Wiener processes with amplitude $\sigma_{ou}$ and time constant $\tau_{ou}$. The parameters used for the WC simulations are listed in Table S2. Equations were integrated using the Heun stochastic method in TVB with a time step of 0.5 ms.

#### 1.2 Supplementary Figures

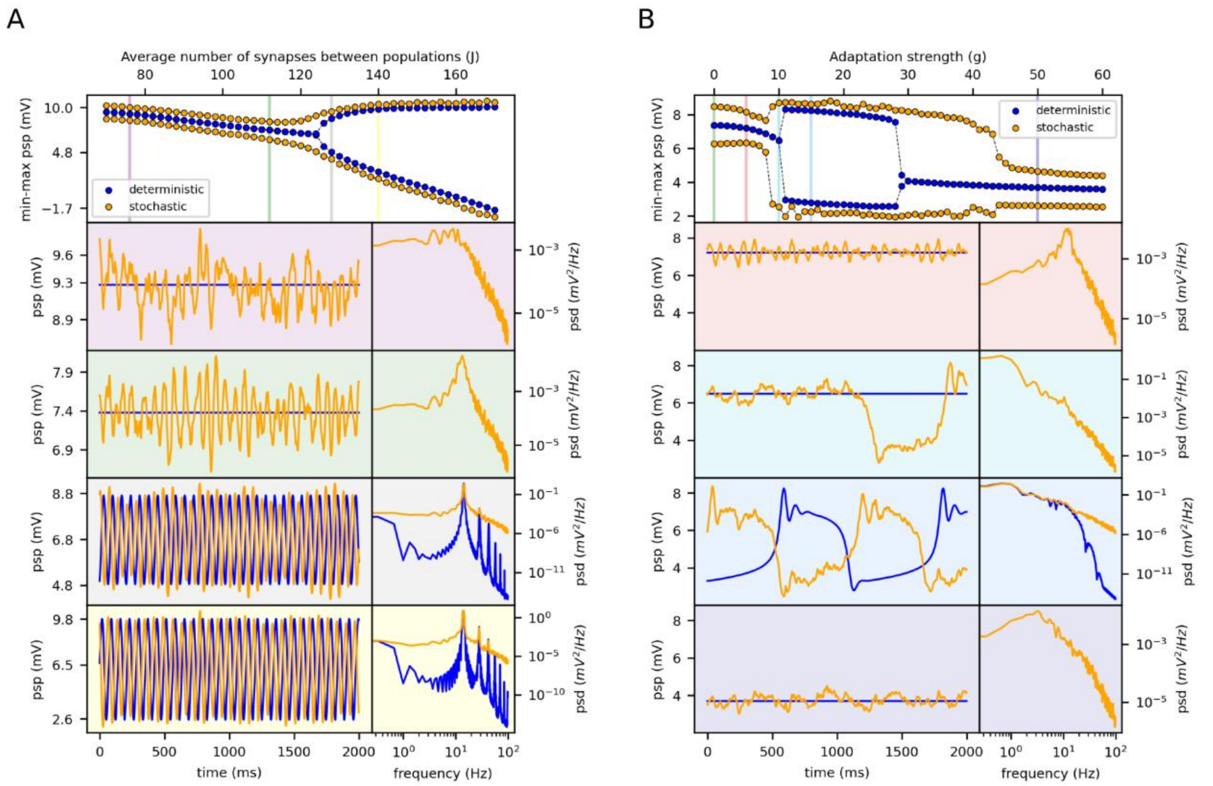


**Figure S1 Bifurcation diagrams and oscillatory dynamics in deterministic and stochastic JR model.** (A) Bifurcation diagram showing the min-max range of postsynaptic potential (PSP) as a function of J, for the deterministic (blue) and stochastic (yellow) models. Vertical bars mark specific values of J, for which representative psp traces (left) and their power spectral densities (psd, right) are shown. The background color of each panel corresponds to the respective vertical colored bar in the bifurcation diagram. The model was run without adaptation (g=0). A supercritical Hopf bifurcation occurs around J=128. (B) Bifurcation diagram as a function of adaptation strength (g), with the same conventions as in (A). The synaptic strength J is set to 112 (matching the green vertical bar in panel A). When g=0 the system corresponds to the same configuration of panel (A) for J=112. For low g, the system is near a supercritical Hopf bifurcation and exhibits damped oscillations. For g>9, Up/Down oscillations emerge. At g=9 and g=10, stochastic fluctuations drive transient transitions to the Down state. For g>39, the system stabilizes in a persistent Down state, with occasional Up state excursions for 30<g<39. Vertical bars indicate specific values of g (i.e., g=0, g=5, g=10, g=15, g=50), for which corresponding PSP traces and psd are shown in the lower panels, color-coded accordingly. Stability analysis of the deterministic model was performed in XPPAUT.

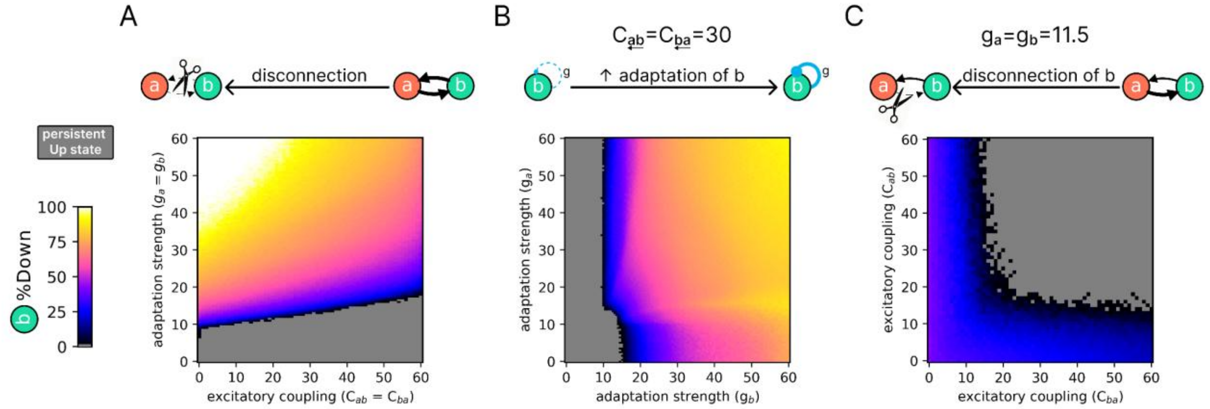


**Figure S2 Effects of adaptation strength and excitatory coupling on slow wave emergence in population $JR_b$.** Each heatmap shows the percentage of time spent in the Down state (%Down) in $JR_b$ as a function of (A) symmetric excitatory coupling ($C_{ab} = C_{ba}$) and adaptation strength ($g_a = g_b$), (B) asymmetric adaptation ($g_a$, $g_b$) with fixed symmetric coupling ($C_{ab} = C_{ba} = 30$), and (C) asymmetric coupling ($C_{ab}$, $C_{ba}$) with fixed adaptation strength ($g_a = g_b = 11.5$). The color scale indicates the proportion of time spent in the Down state, from persistent Up state (gray) to predominant Down state (white).

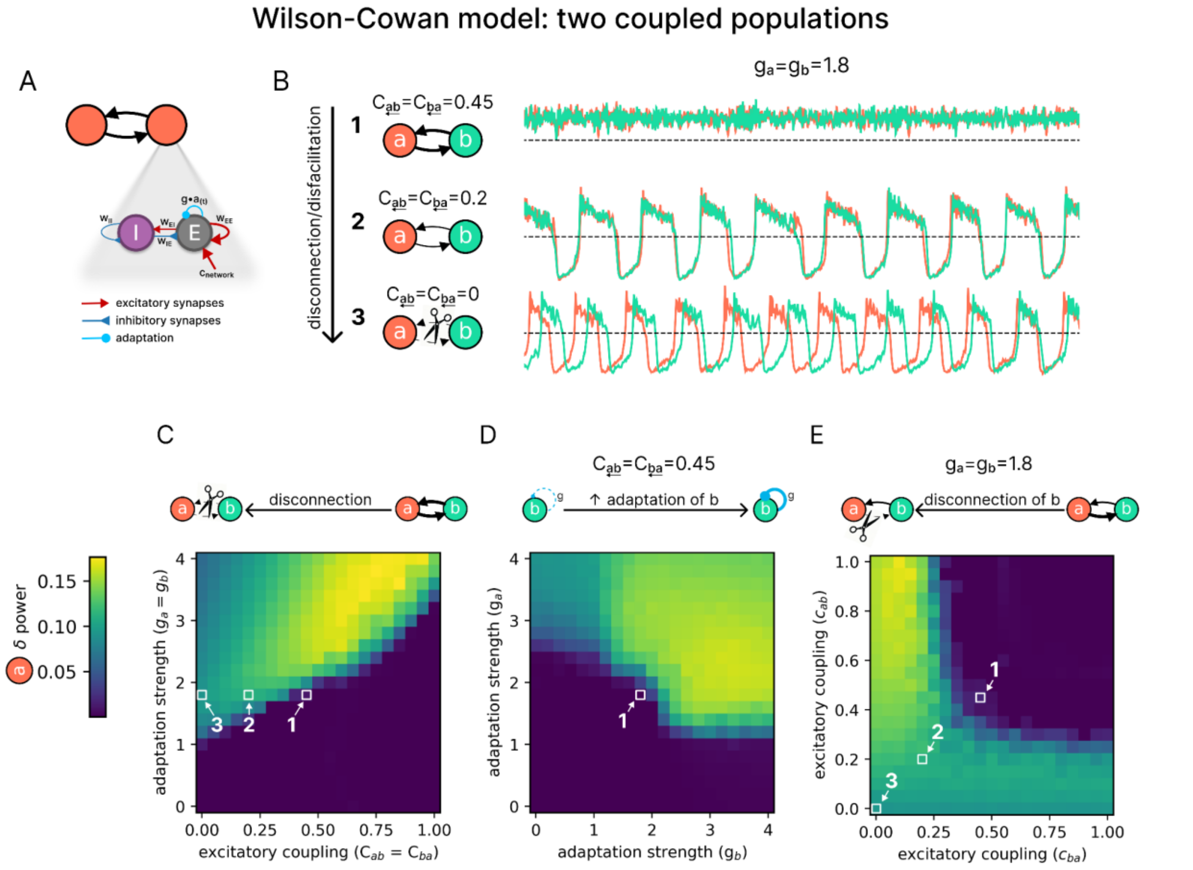


**Figure S3 Effects of adaptation strength and excitatory coupling on slow wave emergence in two coupled WC nodes.** (A) Representation of the Wilson-Cowan (WC) model with spike-frequency adaptation, consisting of interconnected excitatory (E) and inhibitory (I) neuronal pools. (B) Example dynamics of two symmetrically coupled populations ($WC_a$ in orange, $WC_b$ in cyan) for three levels of excitatory coupling ($C_{ab}$=$C_{ba}$). With C=0.45 and g=1.8 (top, #1), both populations show fast oscillations in a persistent Up state. Reducing coupling to C=0.2 (middle, #2) induces a transition to slow oscillations (SO). Complete disconnection (C=0, bottom, #3) further enhances SO activity.. (C) Heatmap showing the δ power of node $WC_a$ as a function of adaptation strength (g, y-axis) and symmetric excitatory coupling (C, x-axis). The three squares with numbers correspond to the configuration of panel B, highlighting the effect of reducing C with g=0.45 kept fixed. Notice how the δ power increases through the transition from a persistent Up state (δ power) to SO (high δ power). (D) Effect of asymmetric adaptation. Heatmap shows δ power in $WC_a$ as a function of $g_b$ (x-axis) and $g_a$ (y-axis), with symmetric coupling fixed at C=0.45. From the square #1, varying $g_b$ (i.e., changing the adaptation of $WC_b$) alone can induce a transition from a persistent Up state to SO (high δ power) in $WC_a$, despite $g_a$ being kept constant. (E) Effect of asymmetric coupling. Heatmap shows δ power in $WC_a$ as a function of $C_{ba}$ (x-axis) and $C_{ab}$ (y-axis), with g=1.8 kept fixed. From the square #1, reducing $C_{ba}$ (i.e., disconnecting $WC_b$) alone is sufficient to induce a transition from a persistent Up state to SO (high δ power) in $WC_a$, even when $C_{ab}$ remains constant. Numbers 1, 2, and 3 in all panels correspond to identical parameter configurations across plots.

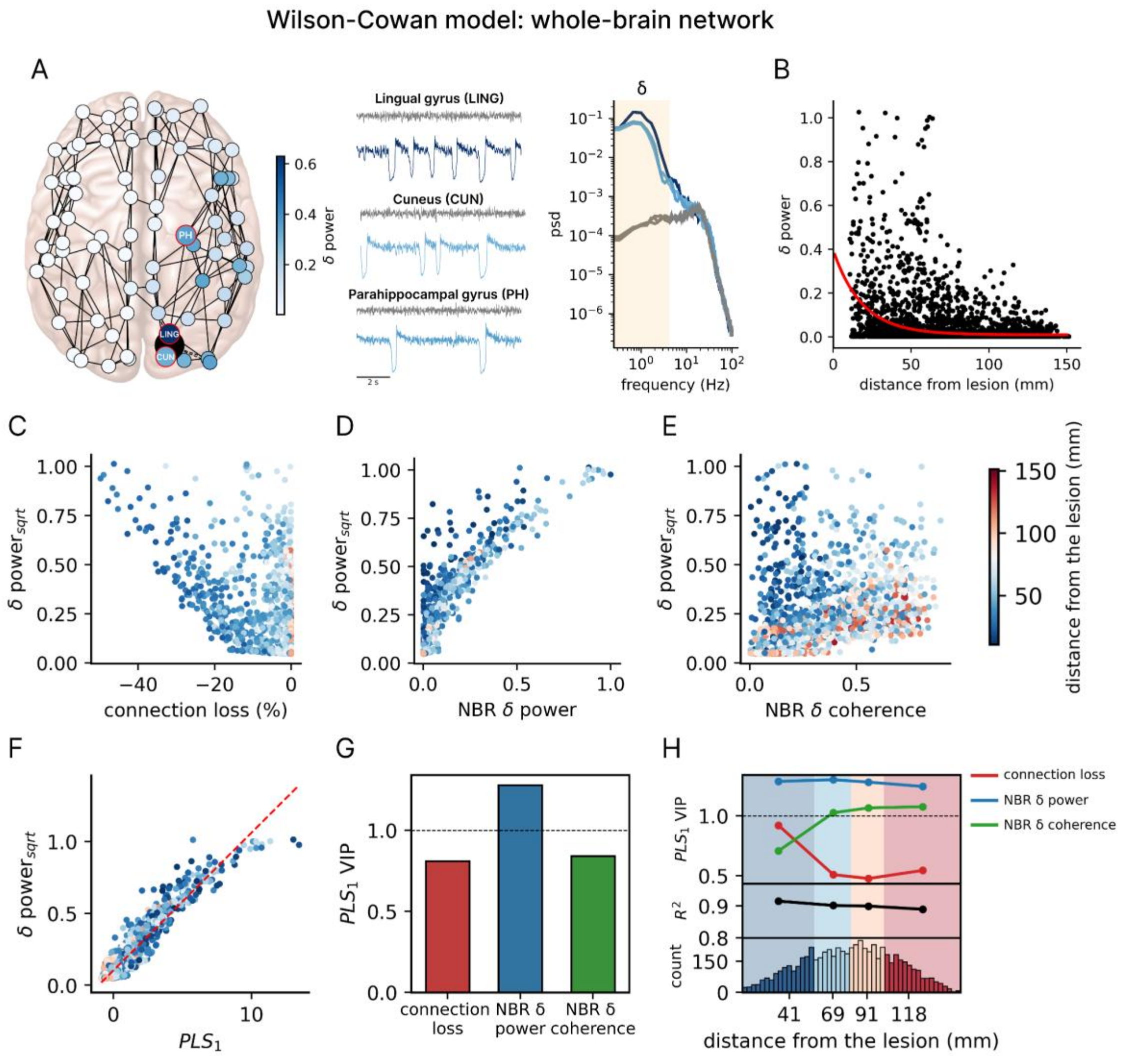


**Figure S4 Slow wave generation and propagation following lesion in a whole-brain network of WC nodes.** (A) Whole-brain network model with nodes defined by the Automatic Anatomical Labeling (AAL) atlas. The lesion of a node (here, the right calcarine cortex) is represented as a black dot, with the impact on the intact nodes quantified with δ power. The node colors indicate δ power, as shown in the accompanying color bar. Traces illustrate simulated activity from three representative regions in the pre-lesion (control; gray) and post-lesion (blue hues) conditions. In the control condition, the nodes display low-amplitude and fast oscillations in a persistent Up state (awake-like regime). After the lesion, SWs emerge along with an increase in δ power in the power spectral density (psd). The δ band is highlighted by the yellow span in the psd. (B) δ power as a function of the Euclidean distance from the lesion. The red curve represents an exponential fit ($R^2$=0.11), indicating a distance-dependent decay in SW propagation. (C) Relationship between connection loss (percentage of lost coupling strength due to the lesion relative to control) and the square root of δ power (δ $power_{sqrt}$). A negative relationship is observed (r=-0.57). (D) Relationship between the δ power of neighboring nodes (NBR δ power) and δ $power_{sqrt}$. A positive relationship is observed (r=0.89). (E) Relationship between the coherence of SW activity of neighboring nodes (NBR δ coherence) and δ $power_{sqrt}$. The variables show a positive relationship (r=0.59). (F) Partial least squares regression (PLS) of δ $power_{sqrt}$ and the score of the first PLS component ($PLS_1$) derived from the three independent variables (connection loss, NBR δ power, and NBR δ coherence). The model explained ~90% of the variance ($R^2$=0.92; after cross-validation $R^2$=0.91). (G) $PLS_1$ weights for the three independent variables, indicating their relative contributions. (H) Distance-dependent contributions of the three independent variables to δ $power_{sqrt}$. Nodes were divided into quartiles based on their distance from the lesion, and PLS regression was performed for each subset. For each quartile, the model's $R^2$ and the VIP scores of each variable are reported.

#### 1.3 Supplementary Tables

**Table S1 Parameters of the Jansen-Rit model used in the simulations.** The table lists the parameters used in The Virtual Brain (TVB) implementation of the Jansen-Rit (JR) model with spike-frequency adaptation. All values lie within physiological ranges and have been previously explored in the literature (e.g., J and $V_0$[3]; A, B, a, and b[4]; $k_{ad}$[5]). The excitatory coupling strength factor c and the adaptation strength g were varied in the study and were set to c=30 and g=11.5 for both the toy and whole-brain networks. The parameter J was explored within the range used in[3] and was set to produce subcritical dynamics just before a supercritical Hopf bifurcation, resulting in a peak in the power spectrum in the alpha band (J=121 for the whole-brain network model and J=112 for all other models).

| Parameter | Description | Value |
|---|---|---|
| A | Maximum amplitude of EPSP. Also called average synaptic gain | 3.25 mV |
| B | Maximum amplitude of IPSP. Also called average synaptic gain | 36.67 mV |
| a | Reciprocal of the time constant of the passive membrane and all other spatially distributed delays in the dendritic network. Also called average synaptic time constant | 0.1 $ms^{-1}$ (=10 ms) |
| b | Reciprocal of the time constant of the passive membrane and all other spatially distributed delays in the dendritic network. Also called average synaptic time constant | 0.083 $ms^{-1}$ (=12 ms) |
| $v_0$ | Firing threshold (PSP) for which a 50% firing rate is achieved | 5.52 mV |
| $v_{max}$ | Determines the maximum firing rate of the neuronal pools | 0.0025 $ms^{-1}$ |
| r | Steepness of the sigmoidal transformation | 0.56 $mV^{-1}$ |
| J | Average number of synapses between the neuronal pools | varied |
| $a_1$ | Average probability of synaptic contacts in the feedback excitatory loop | 1 |
| $a_2$ | Average probability of synaptic contacts in the slow feedback excitatory loop | 0.8 |
| $a_3$ | Average probability of synaptic contacts in the feedback inhibitory loop | 0.25 |
| $a_4$ | Average probability of synaptic contacts in the slow feedback inhibitory loop | 0.25 |
| μ | Mean input firing rate | 0.22 |
| $k_{ad}$ | Adaptation rate constant | 0.001 $ms^{-1}$ |
| g | Adaptation strength | varied |
| $\sigma_{noise}$ | Noise strength. Standard deviation of the Gaussian distribution used to generate the additive white noise | $10^{-4}$ |
| c | Factor scaling the excitatory coupling strength between nodes | varied |
| v | Signal propagation velocity | 4 m/s |

**Table S2 Parameters of the Wilson-Cowan model used in the simulations.** The table lists the parameters used in The Virtual Brain (TVB) implementation of the rate model with spike-frequency adaptation. All parameter values were taken from the original publication[1], except for the time constant of the adaptation dynamics ($tau_a$, changed here from 200 to 500 ms to better align with the JR model while remaining consistent with adaptation kinetics[6]) and the threshold of the inhibitory input-output function ($h_i$ changed from 12 to 9 to increase the contribution of the inhibitory population). In the two-coupled-population model, the adaptation strength (g; denoted as b in the original article) and the excitatory coupling strength (c) were varied, whereas in the whole-brain model, these were fixed to 1.8 and 0.45, respectively, to produce an awake-like regime in the intact model (pre-lesion).

| Parameter | Description | Value |
|---|---|---|
| $\tau_{e/i}$ | Time constant of the excitatory/inhibitory population | 5 ms |
| $\tau_a$ | Time constant of the adaptation dynamics | 500 ms |
| $w_{ee}$ | Internal coupling from excitatory to excitatory neurons | 4 |
| $w_{ei}$ | Internal coupling from excitatory to inhibitory neurons | 4 |
| $w_{ie}$ | Internal coupling from inhibitory to excitatory neurons | 3 |
| $w_{ii}$ | Internal coupling from inhibitory to inhibitory neurons | 0 |
| $k_{e/i}$ | Amplitude or scaling factor for the excitatory/inhibitory input-output function | 0.02/0.05 |
| $h_{e/i}$ | Threshold of excitatory/inhibitory input-output function | 0/9 |
| n | Power law exponent of input-output functions; determines the sharpness of the population response | 2 |
| $a_a$ | Gain of the adaptation dynamics | 5 |
| $mu_a$ | Firing threshold of the adaptation | 1.5 |
| g | Adaptation strength | varied |
| $I_{e/i}$ | External input to excitatory/inhibitory population | 3.6 |
| $\sigma_{ou}$ | Noise strength. Standard deviation of the Ornstein-Uhlenbeck process | 0.002 |
| $\tau_{ou}$ | Correlation time of the Ornstein-Uhlenbeck process | 5 ms |
| c | Factor scaling the excitatory coupling strength between nodes | varied |
| v | Signal propagation velocity | 4 m/s |

#### 1.4 Supplementary References


1. Levenstein, D., Buzsáki, G. & Rinzel, J. NREM sleep in the rodent neocortex and hippocampus reflects excitable dynamics. *Nat Commun* **10**, 2478 (2019).
2. Ahmadian, Y., Rubin, D. B. & Miller, K. D. Analysis of the stabilized supralinear network. *Neural Comput* **25**, 1994–2037 (2013).
3. Jansen, B. H. & Rit, V. G. Electroencephalogram and visual evoked potential generation in a mathematical model of coupled cortical columns. *Biol Cybern* **73**, 357–366 (1995).
4. David, O. & Friston, K. J. A neural mass model for MEG/EEG: coupling and neuronal dynamics. *NeuroImage* **20**, 1743–1755 (2003).
5. Moran, R. J. *et al.* A neural mass model of spectral responses in electrophysiology. *Neuroimage* **37**, 706–720 (2007).
6. Benda, J. & Herz, A. V. M. A universal model for spike-frequency adaptation. *Neural Comput* **15**, 2523–2564 (2003).
